# Three-dimensional molecular architecture of mouse organogenesis

**DOI:** 10.1101/2022.11.17.516228

**Authors:** Fangfang Qu, Wenjia Li, Jian Xu, Ruifang Zhang, Jincan Ke, Xiaodie Ren, Xiaogao Meng, Lexin Qin, Jingna Zhang, Fangru Lu, Xin Zhou, Xi Luo, Zhen Zhang, Guangming Wu, Duanqing Pei, Jiekai Chen, Guizhong Cui, Shengbao Suo, Guangdun Peng

## Abstract

Mammalian embryos have sophisticated cell organizations that are orchestrated by molecular regulation at cellular and tissue level. It has recently been appreciated that the cells that make up the animal body themselves harbor significant heterogeneity in the context of both cellular and particularly spatial dimension. However, current spatial transcriptomics profiling of embryonic tissues either lack three-dimensional representation or are restricted to limited depth and organs. Here, we reported a holistic spatial transcriptome atlas of all major organs at embryonic day 13.5 of mouse embryo and delineated a 3D rendering of the molecular regulation of embryonic patterning. By integrating with corresponding single-cell transcriptome data, the spatial organogenesis atlas provides rich molecular annotation of the dynamic organ nature, spatial cellular interaction, embryonic axes and divergence of cell fates underlying mammalian development, which would pave the way for precise organ-engineering and stem-cell based regenerative medicine.

## Introduction

Organogenesis sets up the functional layout of the animal body from three germ layers, which involves intensive cell-cell interaction, cell fate determination, cell proliferation, as well as spatial arrangement of cells into distinct tissues and ultimately functional organs. Recently, technologies to measure transcriptomes from single cells or leveraging in vitro stem cell-based organoids models have opened up new avenues for understanding organ development ^1^. For example, systematic single-cell atlas reveals hundreds of cell types and states, and illustrates developmental trajectories for many organs at mouse organogenesis ^2^. Similarly, human embryo organogenesis at single-cell resolution has also been reported by characterizing major cell types and developmental regulation programs ^3^. However, being an anatomically stringent organization, organogenesis embryos have high order of tissue architecture and location-dependent mechanisms that are masked by dissociation-based single-cell genomics.

Spatially resolved transcriptome technology represents a significant endeavour to glean unprecedented views of the molecular regionalization in complex tissues and process. Recently, spatial molecular architecture of embryo development has been unraveled by emerging spatial transcriptome approaches such as Geo-seq ^4^, seqFISH ^5^, DBiT-seq ^6^ and Stereo-seq ^7–9^. For example, sagittal sections from mouse embryos spanning E9.5 to E16.5 with one-day intervals were spatially mapped to achieve a global view of organogenesis at cellular resolution ^7^. Similarly, one 2D section of a whole E10 embryo was also spatially charted and anatomic annotation of major tissue regions were defined ^6^. However, cells of the embryo in the context of structure and morphology, are in a three-dimensional environment having characteristic biophysical and biomechanical signals not revealed in a 2D setting, which influence cell functions such as migration, proliferation, interaction, patterning and axes formation. Moreover, some cellular processes of differentiation and morphogenesis occur preferentially in 3D instead of 2D. Therefore, a 3D spatial transcriptome profiling of tissue and organ specific microarchitecture to align serial sections of the embryo into an entire organism is vital for our understanding of embryo organogenesis, which has not been reported.

Embryo organogenesis sets a framework for functional manifestation of organs with the synchronized series of cellular and morphological changes. The early stage of organogenesis (E9.5-E13.5) is particularly of interest for its immediate implication for organoids studies, tissue engineering and dissecting major developmental defects ^2,10^. Additionally, knowledge learned from the co-evolving of various organ and their 3D interaction will be instrumental for designing strategy of in vitro organogenesis ^11–13^. Using a 10x Visium platform which accommodates substantial cell coverage and a high gene-detecting ability, we built an organogenesis spatial atlas composing collective transverse-sections at embryonic day 13.5 (E13.5) mouse embryo to represent almost all organ primordia that have been configured at this stage. The spatial atlas of organogenesis embryo presents a comprehensive view of dynamic cell location, cell-cell communication, spatial heterogeneity and organ architecture formation, providing a molecular basis for our understanding of cellular interaction and allocation in mouse organogenesis.

## Results

### Construction of a spatial transcriptome atlas of embryo organogenesis at 13.5

To generate a comprehensive molecular architecture of the embryo development at E13.5, we performed spatial transcriptomics on individual mouse embryos by applying the 10x Genomics Visium platform. The whole embryo was serially cryo-sectioned into about 1,000 sections along craniocaudal axis at 10 μm thickness. To achieve a comprehensive and concise representation of the anatomic structures and molecular profiling, a total of 10 sections (about every 100 sections apart from each other) in a 3D dimension were collected for spatial transcriptomic analysis (Fig. 1a). RNA-sequencing generated a high quality of spatial molecular map with a median depth of 244.7 million reads per library, 16,418 spots from the whole embryo, a median of 5,668 genes and 22,253 unique molecular identifiers (UMIs) per spot (Extended Data Fig. 1a,b). Reassuringly, we compared two sections from a replicate embryo taken from the similar locations and found that gene expression correlation, cluster identifications and spatial expression of marker genes are consistent, suggesting a high reproducibility of the spatial transcriptome and spatial expression domains among different embryos in our spatial early stage-organogenesis atlas (Extended Data Fig. 1c-o). Meanwhile, the spatial distribution of marker genes were consistent with in situ hybridization (ISH) data taken from Mouse Genome Informatics (MGI) or Allen Brain Atlas (ABA) ^14^ (Extended Data Fig. 1o). We thus focused our analysis on tissue sections from the represented embryo.

**Fig. 1.**
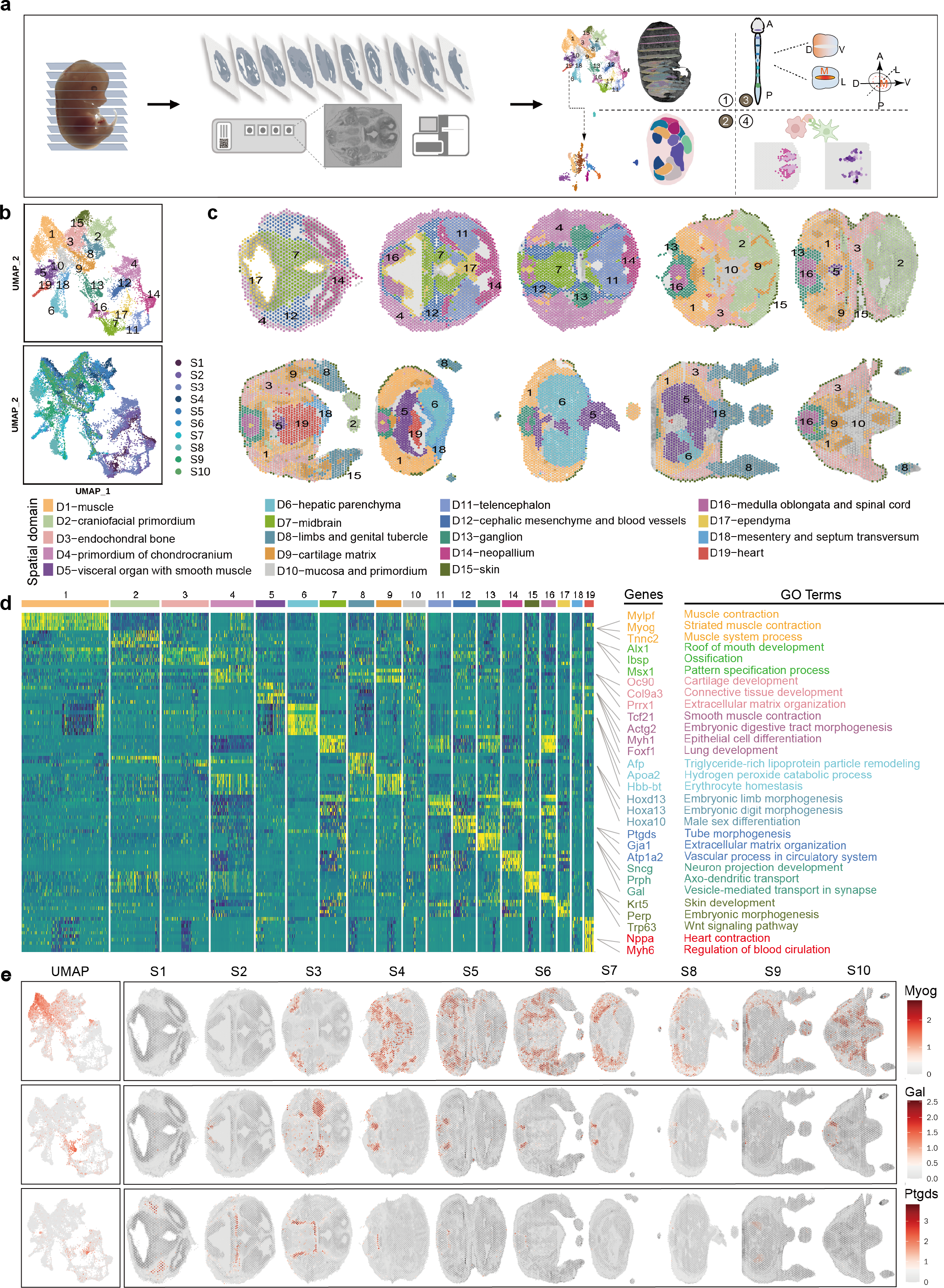
3D spatial transcriptional atlas for mouse organogenesis at E13.5. a. Schematic overview of experimental design and analysis workflow for spatial transcriptome of mouse organogenesis at E13.5 b. UMAP projection and clustering of the spatial transcriptome from all sections and spots, colored by spatial domains (Top) and sections (Bottom). c. The spatial distribution of 19 UMAP clusters in (a) across all embryo tissue sections, annotated according to anatomical structures and molecular features in (d). d. The heatmap showing the expression pattern of the top five representative marker genes for each spatial domain. Examples of marker genes are listed together with the enriched Gene Ontology (GO) terms for selected spatial domains. e. UMAP projection and spatial expression distribution of selected marker genes, *Myog*, *Gal*, and *Ptgds* across all the ten embryo sections. S, section; D, domain.

To systematically reveal the spatial molecular architecture of the E13.5 mouse embryo in the fully represented view, we merged all the 10 sections of the whole embryo and performed dimensional reduction by principal component analysis (PCA) and unsupervised clustering. On the uniform manifold approximation and projection (UMAP), the spatial clusters were separated into two major groups: head and body parts (Fig. 1b). We identified 19 consensus spot clusters and annotated them based on the expression of the signature genes and enriched GO terms (Fig. 1b-d, Extended Data Fig. 2a,b and Supplementary Table 1). The spatial transcriptome atlas covered the majority of tissue or organs at this stage such as brain cerebrum, spinal cord, respiration tract systems, gastrointestinal tract systems, circulation system, bone, skin, gonad and etc. We found the spatial clusters matched well with their defined spatial anatomy structures after mapping spots back to the original coordinates of the tissue sections (Fig. 1c and Extended Data Fig. 2b). The D1-muscle domain (marked by *Myog* and *Tnnc2*, Fig. 1e and Extended Data Fig. 2a,b) were shared across all 7 sections of the body parts, while brain related clusters such as domain D7-midbrain, D11-telencephalon, D14-neopallium and D12-cephalic mesenchyme and blood vessels (marked by *Ptgds and Atp1a2*, Fig. 1e and Extended Data Fig. 2a,b) were distributed in the head parts. Meanwhile, spatial domain D17-ependyma, D16-medulla oblongata and spinal cord, D13-ganglion (marked by *Gal and Sncg*, Fig. 1e and Extended Data Fig. 2a,b) and D9-cartilage matrix were presented across the assayed embryo sections due to a ubiquitous existence (Fig. 1b,c and Extended Data Fig. 2b,c). In addition, we also identified organ-specific clusters. For example, D6-hepatic parenchyma, specifically expressing *Afb and Apoa2* ^15^, is predominately located in liver; D19-heart exhibited differential expression of *Nppa* and *Myh6*, which is associated with heart contraction and blood circulation regulation (Fig. 1d and Extended Data Fig. 2a,b) ^16,17^. Altogether, the organism-level of spatial atlas provided a holistic molecular annotation for tissue architecture of mouse organogenesis at E13.5.

### Regionalization and orchestration of gene regulatory network activity for organ development

To reveal the regionalized transcriptional regulatory activity underlying the spatial gene expression patterns, we implemented the single-cell regulatory network inference and clustering (SCENIC) pipeline ^18^ and calculated the regulon activity score (RAS) for each spatial spot. The spatial domains based on RAS were consistent with the spatial gene expression clusters (Extended Data Fig. 3a-c), indicating the activity of transcription factors (TFs) regulatory network may be involved as the driving force to determine the cell fates and locations.

We then sought to systematically identify critical and functional relevant TF regulators that associate with each spatial domain. By computing the regulon specificity score (RSS) of each regulon for all 19 spatial domains based on Jensen-Shannon divergence ^19^, we identified significantly enriched regulons in each spatial domain (Fig. 2a,b and Supplementary Table 2). Through this, we obtained the organism-level key TF networks specifically function in both location and cell-type dependent manner (Fig. 2a,b and Extended Data Fig. 3d). For example, Hnf1a, Nr1h3 and Cebpe regulons were identified as the top regulons in D6-hepatic parenchyma, in agreement with their functions in hepatocyte development, differentiation or bile acid homeostasis ^20–23^. Bhlhe40 regulon showed specific and strong activity in the D9-cartilage domain and Trp63 exhibited a specific activity in the D15-skin domain. In pan-muscle cells, Myf6, and Myod1, were identified as master TF networks ^24–26^. Of note, we also found that Isl2 and Prrxl1 were shown as the top ranked regulators in D13-ganglion region. Meanwhile, Dlx4 networks exhibited high specificity score in the domain of craniofacial primordium, corroborating their reported roles in regulation of craniofacial bones development ^27^. Importantly, besides known cell type markers, the 3D spatial atlas delineated unappreciated TFs that are enriched in respective organs, for example, Rxrg in muscle cells and Mecom in limb tissues (Fig. 2a,b) which was recently reported for shaping the limbs and digits ^28^. These results suggest that the prioritized unrecognized spatial-domain specific regulons identified from our spatial atlas could serve as an important resource for further function analyses. Taken together, the 3D spatial atlas data unveiled the specifically regionalized transcriptional regulatory network underlying the tissue organization.

**Fig. 2.**
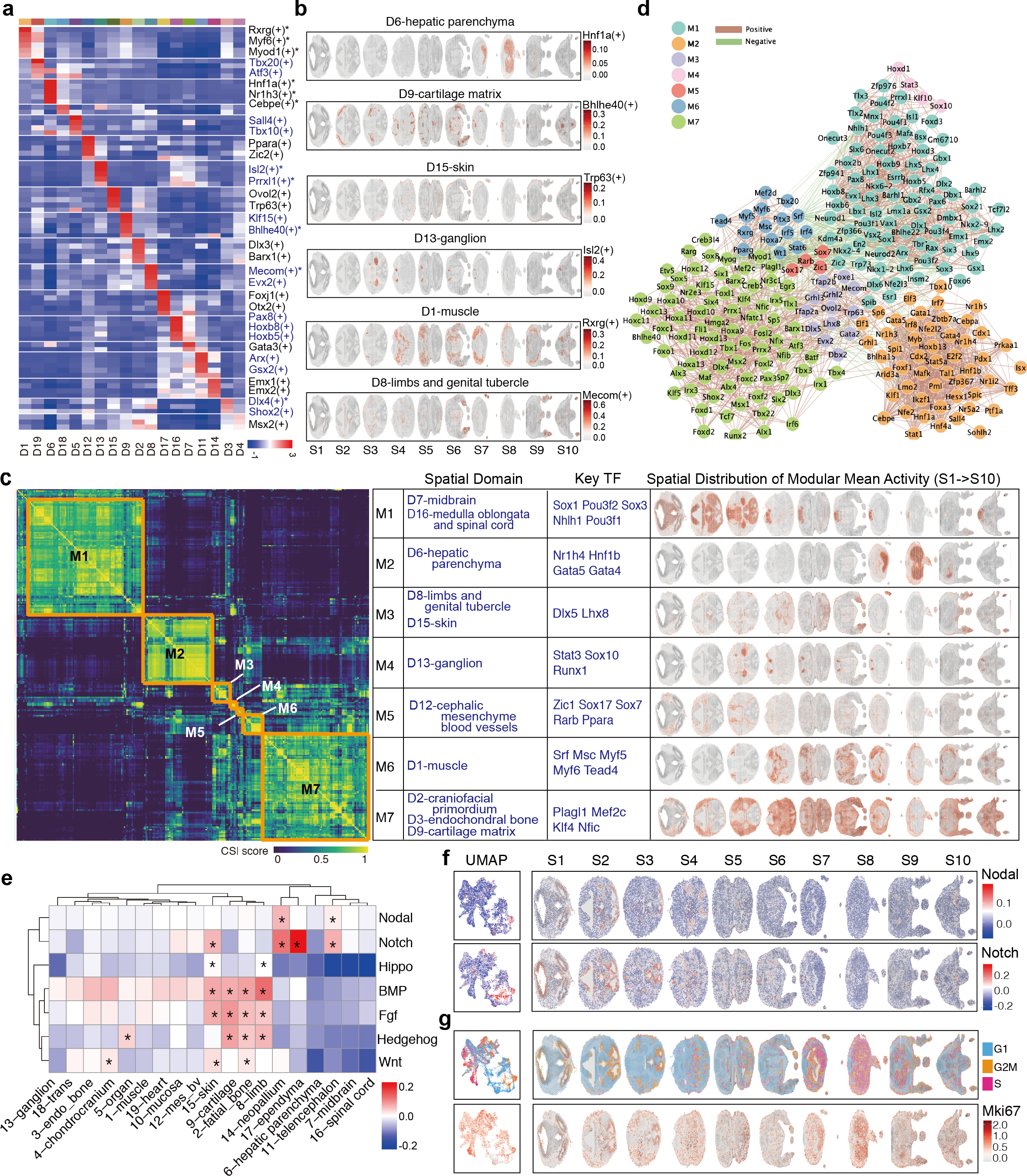
Spatial gene regulation network, signaling and proliferation activity. a. Heatmap of mean regulon activity score of top 5 regional specific regulons for each spatial domain. b. Spatial distribution of regulon activity scores across embryonic tissues for selected domain specific regulons. c. The hierarchical clustering of heatmap showing the seven regulon groups based on CSI matrix, with associated spatial domains, representative TFs and spatial plots visualizing the spatial distribution of mean activity score across each section. d. The co-expression network based on CSI for the seven regulon groups. The node with different colors represents the regulon in each module and width of edge represents the CSI value of two nodes (filtered with CSI >0.85). Color of edges represent positive (brown) or negative (green) correlation. e. Activity of development-related signaling pathways in each spatial domain. Differentially activated signaling in spatial domains computed by Wilcox test are marked with * for mean score greater than 0 and p-value less than 0.001. f. UMAP and spatial plots showing activities of Notch and Nodal signaling in all spots and all sections of embryo tissue. g. Spatial plots showing the spatial distribution of cell cycle activity of G1, G2M and S phase in embryo tissue and the spatial expression of *Mki67*.

During embryogenesis, transcription factors often regulate gene expression in a coordinate and combinatorial manner. The divergent or convergent TF regulation mechanisms outside or inside developmental lineages are particularly interesting. To further reveal the orchestrated regionalized transcriptional regulation networks in our 3D spatial atlas, we calculated the similarity of top selected regulons based on Connection Specificity Index (CSI) ^29^ and applied hierarchical clustering to identify potential function-related regulon groups. The TF regulons formed 7 co-activation modules. By computing the averaged RAS score for each TF regulon module, we found distinct spatial regulon modules, which highlighted the coherence of TF regulons in particular tissue regions (Fig. 2c). To reveal the underlying connections in each module, we further constructed a TF co-expression network on the basis of their CSI value (Fig. 2d). Among them, regulon module 1 (M1) with key regulators such as Sox1 and Pou3f2 showed high activity in D7-midbrain, D16-spinal cord, and cerebrum domains, which indicates general neural development regulation mechanisms. M2 was mainly enriched in hepatic parenchyma, with function involved in myeloid cell differentiation, erythrocyte differentiation, indicating hematogenesis in fetal liver. Regulons within M2 such as *Nr1h4 and Hnf1b* were strongly connected with each other but with lower connection to regulons out of M2 (Fig. 2c,d). For M3 (D8-limbs, D15-skin), M6 (D1-muscle) and M7 (D2-craniofacial primordium, D3-endochondral bone, D9-cartilage matrix) which are related to mesoderm and ectoderm lineages, they have strong positive correlations with each other and shared common regulators (Fig. 2c,d), indicating that cell types in these two lineages may co-develop with each other. Collectively, the regulation network analysis showed regionalized specific regulation and orchestration, which will help to spatially dissect the regulatory mechanism across different cell types and locations in controlling embryo organogenesis.

### Spatial signaling pathway activity in the whole embryo

The patterning and regionalization of embryos during development is highly dependent on morphogenetic signals. In order to systematically chart the spatial layout of the activities of signaling pathways in the whole embryo, we examined the enrichment score of signaling genes (including ligands, receptors, key signaling effectors and regulators) of Wnt, Bmp, Fgf, Hippo, Nodal, Notch and Hedgehog pathways. The comprehensive map of gene expression and spatial domains allow in-depth analysis of mouse mid-gestation organogenesis development both globally and in individual spatial domains (Fig. 2e,f). Of note, the spatial distribution of the signaling scores illustrated overall relatively high activity of developmental signaling across the telencephalon, ependyma, limb, cartilage and skin, whereas especially low activity in liver domain, midbrain and spinal cord, suggesting the differential induction and patterning scheme in different tissue and organs. As an example, Nodal signaling mainly had strong activity in regions of cerebrum, part of the midbrain, and jaw area of the craniofacial primordium, indicating its important roles in neurogenesis and determination of the function of the mouth^30^ (Fig. 2e,f). Meanwhile, the activity of Notch signaling which mediates juxtracrine cell-cell communication ^31,32^, were also enriched in the brain and spinal cord ependyma region, cerebrum and part of the skin domain. Bmp activity showed relative high activity in limb and less represented in liver and cerebrum and midbrain domain, consisting with previous studies on regulation of Bmp signaling during embryogenesis ^33,34^ (Fig. 2e). Whereas, Hedgehog and Fgf signaling also showed moderate activity in cartilage and limb, indicating these two signaling pathways may coordinate with BMP to play important roles in limb and bone morphogenesis (Fig. 2e).

Next, we examined the spatial distribution of the proliferation state by calculating the “Cell cycle score”. Notably, the nervous system including D14-cerebrum, part of D11-cerebrum and D17-ependyma regions revealed especially high level of G2M and S phase score, whereas the midbrain, spinal cord and the ganglion domains showed low activity score for both G2M and S phase, suggesting the locations of neural progenitors are associated with high proliferation ability (Fig. 2g). Interestingly, hepatic parenchyma domain and gonad region of visceral organ domain illustrated high level of G2M and S phase score, and the expression levels of *Mki67* were also high in the regions with high G2M score, demonstrating that the cells in these regions are undergoing rapidly self-renewing (Fig. 2g). Taken together, this analysis uncovered the regionalization of proliferation activity in the whole embryo.

### Spatially resolved molecular characterization of the visceral organs

In the organism-level classification, the spatial domain of D5 (Fig. 1c) was a major cluster consisting of spots from multiple visceral organs. To achieve a finer annotation, spots of D5 were subjected to further unsupervised clustering and we obtained 10 sub-clusters (Fig. 3a,b). According to the distinct spatial expression patterns, the specific expressed marker genes and the top enriched GO terms, we assigned the 10 spatial domains as pancreas, bladder, gonad, stomach, gut, metanephros, lung, umbilical, trachea and mesenchyme (Trachea_Mes) and mesonephros-spleen-superior recess of omental bursa (Nep_sp_Om). The spatial data hence provides a full spectrum of all major visceral organs in the mid-organogenesis embryo (Fig. 3b,c and Extended Data Fig. 4a-c), and we were able to define visceral organ-specific signatures (Supplementary Table 3). For example, *Nkx2-1*, a key factor in regulating the development of brain and lung structures, showed specific expression in both lung and brain ^35^ (Fig. 3d). Stomach was distinguished by the expression of tissue specific gene *Barx1* (Fig. 3d), which has been shown in the control of thoracic foregut specification and tracheo-esophageal septation ^36^. In addition, less-studied genes such as *Fst*, *Tmem200a*, *Egln3* were co-expressed with *Barx1* in this domain, indicating their potential role in stomach development (Extended Data Fig. 4b).

**Fig. 3.**
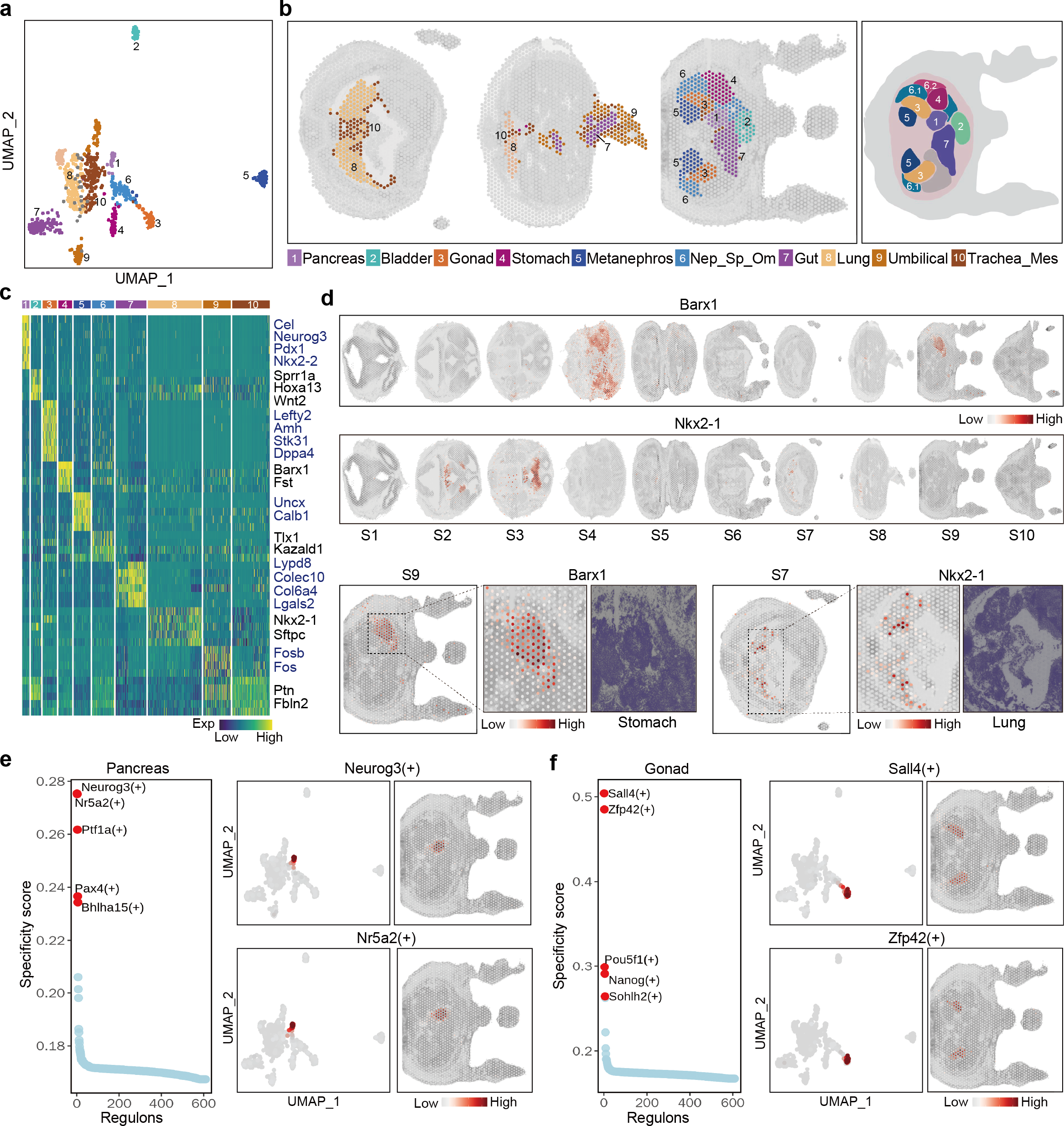
Domain specific gene expression and gene regulation for visceral organs. a. UMAP embedding of spots from D5-visceral organ labeled by subclusters. b. Spatial distribution and annotation of the 10 subclusters of D5-visceral organ. Segmented regions are highlighted on section 9 and colored according to the denoted anatomical structure (right). Subcluster 6 was further divided into 6.1 and 6.2 to represent finer structures. Trachea_Mes, trachea and mesenchyme; Nep_sp_Om, mesonephros-spleen-superior recess of omental bursa. c. The heatmap showing expression pattern of selected top marker genes for each subcluster. d. Spatial expression of selected marker genes *Barx1* for Stomach (top) and *Nkx2-1* for lung (middle) across all embryo sections. The dashed box showing the region-specific expression of *Barx1* in the stomach region of section 9 (bottom left) and *Nkx2-1* (bottom right) in the lung region of section 7 and their corresponding HE-stained tissue regions. e. Rank plot for regulons in pancreas based on regulon specificity score (left). UMAP visualization and spatial distribution of regulon activity scores for the top two regulon Neurog3 (top right) and Nr5a2 (bottom right) in the tissue section 9. f. Rank plot for regulons in gonad based on regulon specificity score (left). UMAP visualization and spatial distribution of regulon activity scores for the top regulon Sall4 (top right) and Zfp42 (bottom right) in the tissue section 9.

Having the high sensitivity and comprehensive tissue coverage, the spatial atlas allows us to exploit signature genes of other embryonic visceral organs which have not been adequately investigated. For example, we found *Sprr1a* showed spatially exclusive expression in bladder tissue (Extended Data Fig. 4b). *Uncx*, *Calb1* and *Foxd1* showed spatially restricted expression in the metanephros regions. *Lypd8, Colec10, Col6a4* were specifically expressed in gut. Among these subdomains, gonad and pancreas expressed more region-specific genes, indicating a strong and unique organ feature (Extended Data Fig. 4a). We identified *Cel, Neurog3, Nkx2-2*, and *Pdx1* to be markers for pancreas development, and *Lefty2, Tex19.1 and Dppa3* to be regional specific genes for Gonad (Fig. 3c and Extended Data Fig. 4b). These findings therefore showcased the valuable utilities of 3D spatial atlas in dissecting sophisticated tissue organizations. Accordingly, we also identified regional specific TF regulons in distinguishing the subregions of these visceral organs (Supplementary Table 4). For example, Neurog3, Nr5a2, Ptf1a, Pax4, and Bhlha15 are associated with pancreas development as also indicated previously ^37–39^ (Fig. 3e), whereas Sall4, Zfp42, Pou5f1, Nanog and Sohlh2 are the highly expressed regulons associated with Gonad (Fig. 3f) ^40^. Nkx2-1 and Cdx2 are the key regulons for lung and Gut respectively (Extended Data Fig. 4d). Our data therefore characterized the previously transcriptionally unappreciated tissue structures within these subclusters of visceral organs.

### The craniocaudal, dorsoventral and radial axes in establishing the spatial patterning of spinal cord

As a prominent developmental event at E13.5, neural tube acquires craniocaudal, dorsoventral and radial positional identities to form brain and spinal cord system. Previously, single-cell RNA-seq has provided many insights into the nervous system development ^41^, and now with the 3D spatial transcriptome, we sought to explore the spatial patterning of spinal cord at the whole organism level.

Homobox (Hox) genes are well known in regulating boundary positioning of distinct neuronal subtypes within both hindbrain and spinal cord along anterior-posterior (AP, craniocaudal) axis, forming the Hox code to establish the regional identities ^42^. We first examined the expression of a total of 26 expressed Hox genes in the spatial domain of D16−medulla oblongata and spinal cord (Fig. 4a,b). Hox genes are generally divided into anterior (Hox 1-3/4), trunk/central (Hox 4/5–9) and posterior (Hox 10-13) paralogues groups (PGs), reflecting their arrangements along each genome cluster ^43^. Indeed, the anterior Hox PGs were mainly expressed in the rostral sections and the posterior Hox PGs were increased in the posterior regions (Fig. 4a). In order to identify novel A-P patterning genes, we firstly arranged the spots along anterior to posterior plane ordered by the expressed Hox genes to establish a pseudo-space trajectory (Fig. 4b), and then identified distinct A-P patterning candidates through differential gene analysis according to the predicted pseudo-space ordering and further applied correlation analysis between gene expression of spots and combinatorial pattern of AP slices to select top related A-P patterning genes (Extended Data Fig. 5a,b and Supplementary Table 5). We therefore identified novel A-P regional specific genes including *Igfbpl1* (Fig. 4c) and *Pou4f1* (Extended Data Fig. 5b) in the rostral region, *Atp1a3* in the central and *AC162693.1* in the caudal region (Fig. 4c), which expand the pool of potential regulators of body plan.

**Fig. 4.**
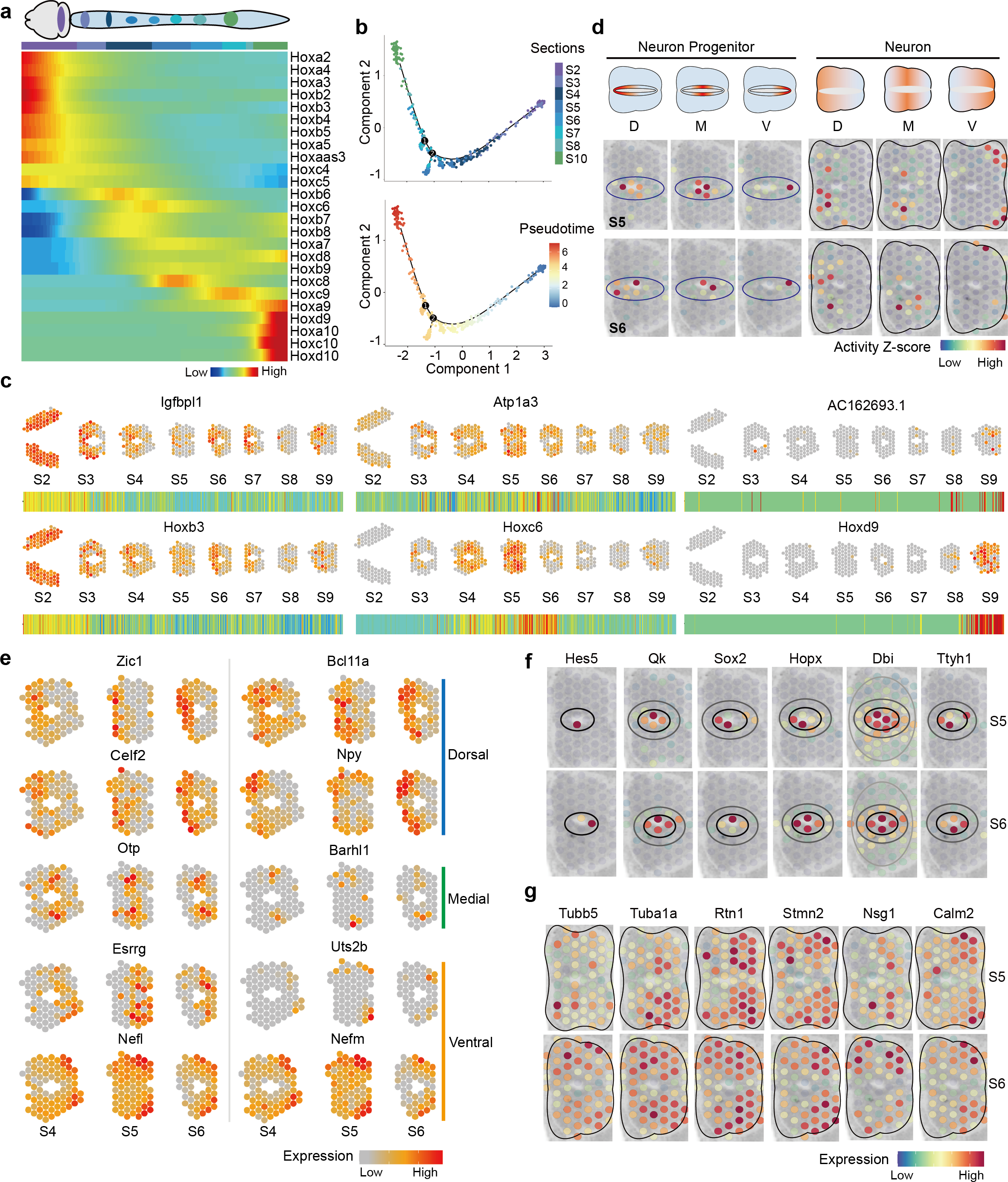
3D alignment to reveal body axes and spinal cord patterning. a. Heatmap plot showing the smoothed spatial expression pattern of Hox family genes along the A-P axis with spots from hindbrain and spinal cord regions ordered by pseudo-axis within each section. b. Pseudo-time trajectory plot showing pseudo-space patterning of spots from hindbrain and spinal cord region across sections of whole mouse embryos, coloured by section numbers (top) and pseudo-time (bottom) . c. Spatial expression of selected Hox family genes and newly identified A-P axis related genes in hindbrain and spinal cord from sections along anterior to posterior (top), and heatmap showing the expression pattern of corresponding genes ordered along sections from anterior to posterior (bottom). d. Spatial plot showing the D-M-V activity scores for respective neuronal progenitors and neuronal cells in spinal cord of section 5 and 6. e. Spatial expression of selected D-V axis related genes in sections 4, 5 and 6. f. Spatial visualization of neural progenitor specific genes *Hes5*, *Qk*, *Sox2*, *Hopx*, *Dbi* and *Ttyh1* in the spinal cord region from section 5 and 6. g. Spatial visualization of neuron marker genes *Tubb5*, *Tuba1a*, *Rtn1*, *Stmn2*, *Nsg1* and *Calm2* in spinal cord region from section 5 and 6.

Next, we took advantage of our transverse collection strategy to examine the dorsal-ventral (D-V) patterning for embryonic spinal cord development. During the development of embryonic spinal cord, structurally and functionally diverse neurons are generated along the D-V axis under the control of precise spatial expression genes and signals ^44,45^. We first allocated the combinatorial expression of a curated marker gene list that are used to define different domains of progenitors ^41^ to the spinal cord spots, including dorsal gene sets (D) by combinatorial marker expression of roof plate (RP) and dp1-dp6, medial gene sets (M) by marker territory of p0-p2, as well as ventral (V) gene sets by pMN, p3 and floor plate (FP) genes. Similarly, we classified gene sets for neuronal regions into Dorsal (D, marked by dl1-dl6), Medial (M, marked by V0-V2b) and Ventral (V, marked by Mn and V3) subdomains. We found well separation of dorsal, medial and ventral structures in the spinal cord regions (Fig. 4d). The activity of dorsal neuronal progenitor markers was predominantly high in the dorsal spots of ventricular zone (Fig. 4d left). Similarly, corresponding neuronal cell type markers were exhibited in an alignment of dorsal to ventral distribution (Fig. 4d right). To predict novel patterning genes in the spinal cord, we identified domain-specific genes along the D-V axis in spinal cord regions (Fig. 4e, Extended Data Fig. 5c and Supplementary Table 6). We found that *Zic1*, *Bcl11a*, *Celf2* and *Npy* were specifically expressed in the dorsal region; *Otp* and *Barhl1* were up-regulated in the medial region; *Esrrg*, *Uts2b, Nefl* and *Nefm* were highly expressed in the ventral region (Fig. 4e). Interestingly, Pou4f1 also exhibit specific expression in the dorsal region, besides its rostral expression along the craniocaudal axis. We verified the expression of *Pou4f1* in the anterior and dorsal regions of neural tube by in situ hybridization (ISH) (Extended Data Fig. 5b).

To further identify the spatial differences in the radial axis (medial-lateral) which is related to neural progenitor proliferation and differentiation, we conducted differentially expressed gene (DEG) analysis along the inner and outer layers for spinal cord spots (Fig. 4f,g). We found well-known marker genes of neural progenitors such as *Hes5*, *Qk* and *Sox2*^46–49^, as well as genes with limited study in neural progenitor cells, such as *Hopx*, *Dbi* and *Ttyh1* which were expressed in the inner regions; whereas genes including *Tubb5*, *Tuba1a*, *Rtn1*, *Stmn2*, *Nsg1* and *Calm2* which may be related to differentiated neuronal cells were highly expressed in the outer layers. We confirmed the locality of *Ttyh1* expression by ISH (Extended Data Fig. 5d). Our analysis therefore suggests that many more spatially restricted genes can be identified to constitute the regionalized patterning of neural tube, hence the spatial atlas may provide new clues for neuronal specification and patterning of developing spinal cord.

### Integrating of single cell and spatial atlas illustrates the spatially resolved cell interaction

Each spot in Visium platform is supposed to be represented by a mixture of cells (~20 cells) and probably consist of several cell types. To investigate the cell type heterogeneity in the spatial regions, we performed cell type deconvolution by Robust Cell Type Decomposition (RCTD) ^50^ with E13.5 mouse single cells derived from TOME dataset as the reference ^1^. We defined specific spatial distribution for 46 cell types in each spatial domain at E13.5 stage (Fig. 5a, Extended Data Fig. 6, 7 and 8 and Supplementary Table 7). As expected, myocytes exhibited the highest abundance in spatial and endothelial cells and white blood cells were also ubiquitously distributed (Fig. 5a and Extended Data Fig. 6a).

**Fig. 5.**
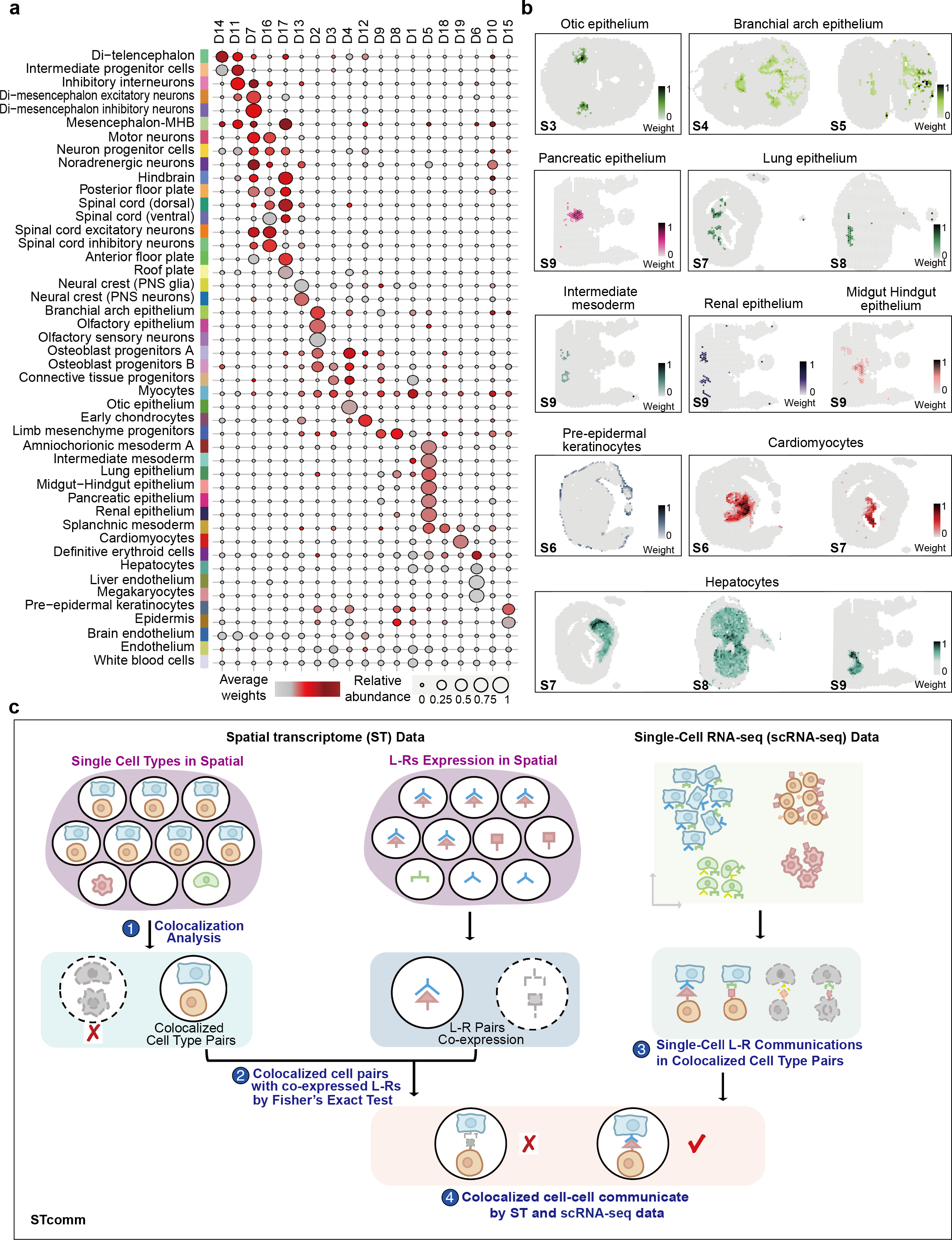
Spatial mapping of cell populations across all the tissue sections. a. Deconvolution analysis inferred the weights of 46 cell populations in all spots of embryo tissue. The dot plot showing cell type composition within each spatial domain. Colour bar indicates the averaged cell-type weights in each spatial domain. Dot size represents the relative abundance of cell types in each spatial domain. b. Spatial distribution of organ/tissue specific cells including epithelium, intermediate mesoderm, pre-epidermal keratinocytes, cardiomyocytes and hepatocytes. c. The workflow of STcomm analysis pipeline which combined the spatial cellular colocalization and L-R co-expression from spatial transcriptomic data with cell-cell communication inferred from sc-RNA seq data (See Methods).

Visualization of the weights of deconvoluted cell types on the tissues showed that the assigned coordinate of organ specific cells corroborates known tissue function (Extended Data Fig. 6, 7 and 8), such as epithelium cells, which were located as single cell types of optic epithelium, branchial arch epithelium, lung epithelium, pancreatic epithelium and renal epithelium (Fig. 5b). By examining the top abundant cell types in each spatial domain (Extended Data Fig. 8b), we found a consensus between cell types and their anatomical structure. For example, D6-hepatic parenchyma was dominated by hepatocytes and definitive erythroid lineage (Fig. 5b and Extended Data Fig. 8b). Similarly, the dominant cell types in D19-heart were cardiomyocytes and endothelium which are essential functional units of heart (Fig. 5a,b and Extended Data Fig. 8b). Interestingly, the early chondrocytes, osteoblast progenitors A, osteoblast progenitors B, and connective tissue progenitors, were located in the different but physically proximity area (Extended Data Fig. 6b), indicating their potentially close interactions. Of note, posterior floor plate cells were detected within a limited area near the ventral region of spinal cord, with only one to two spots for sections across the body trunk, indicative of a high fidelity of the spatial mapping (Extended Data Fig. 7b). The identified marker genes in TOME dataset such as *Slit1*, *Ntn1* and *Shh* showed specific spatial expression in the corresponding spatial regions of posterior floor plate ^1^ (Extended Data Fig. 7b).

To investigate how spatial proximity of cell types may influence each other and shape the signaling landscape to coordinate the developmental programs, we developed a spatial cell-cell communication (CCC) analysis workflow (named STcomm), which combined the spatial cellular colocalizations with their enriched ligand-receptor (L-R) co-expression patterns inferred from both spatial and single-cell transcriptomic data under the assumption that spatially co-localized cells in spot level can more reliably infer CCC mediated by LRs (Fig. 5c and Methods). Firstly, we quantified the colocalization of cell type pairs within spots by calculating Pearson correlation coefficient based on cell type composition predicted by RCTD. We thus identified significant co-occurrence cell type groups belonging to the same spot from the cell type colocalization network (Extended Data Fig. 9a). For example, Olfactory epithelium and Olfactory sensory neurons were co-localized in craniofacial primordium at the front region of section 4 (Fig. 6a). Pre-epidermal keratinocytes and epidermis significantly co-existed and mainly distributed in D15-skin, which are in agreement with expected cell type localization (Extended Data Fig. 9a). Whereas hepatocytes, definitive erythroid lineage, megakaryocytes and liver endothelium formed a colocalization network, indicative of the cellular composition in liver microenvironment. Of note, we found neuron progenitor cells showed spatial proximity with inhibitory interneurons, hindbrain, Di-mesencephalan inhibitory neurons, Di-mesencephalan excitatory neurons and spinal cord excitatory and inhibitory neurons. Neural crest -PNS neurons and neural crest-PNS glia which were abundant in D13-ganglion domain (Fig. 5a) were significantly interacted with each other (Extended Data Fig. 9a) .

**Fig. 6.**
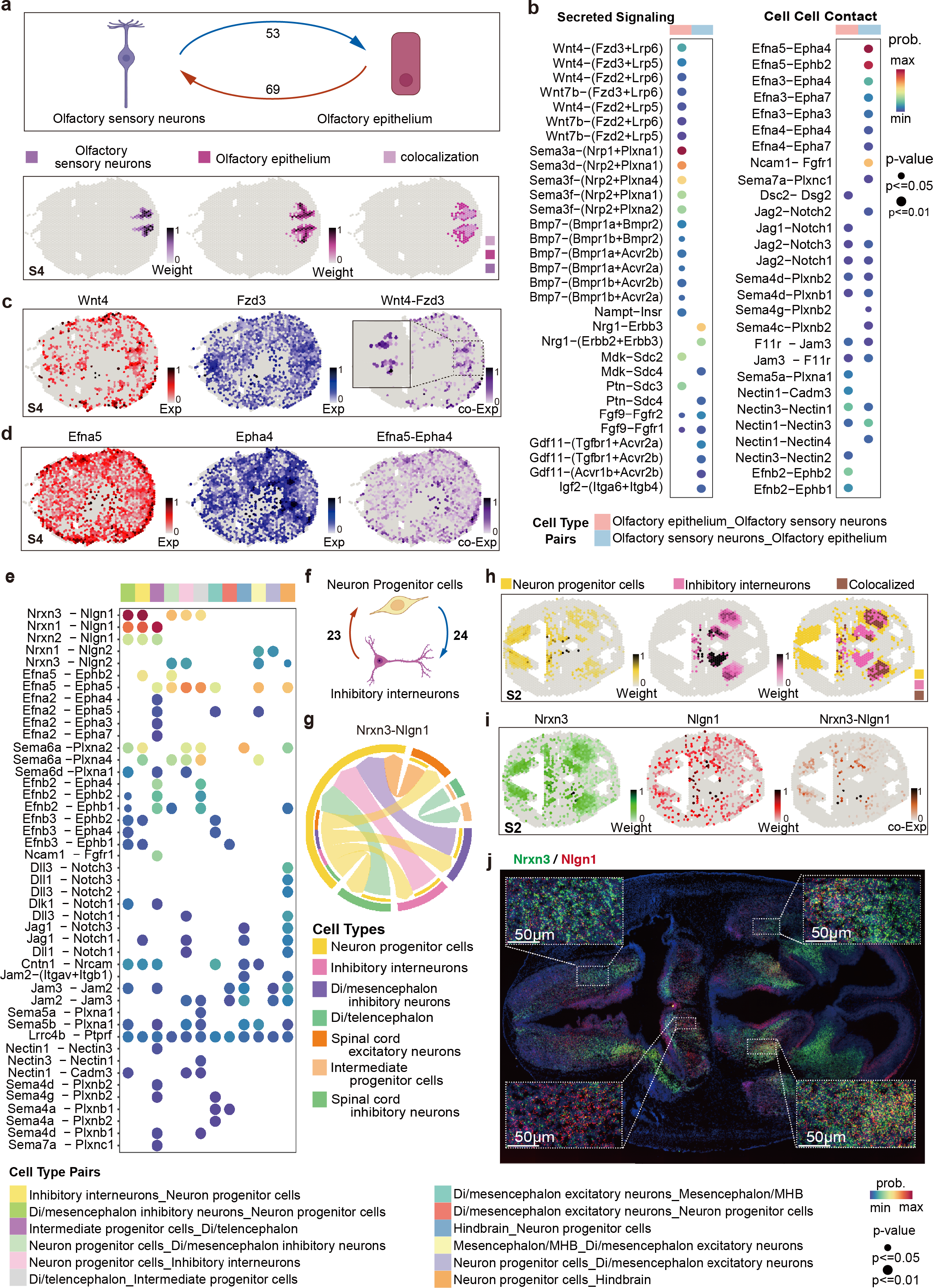
Cell–cell communication network of spatial proximity brain related cell types in mouse embryo organogenesis. a. Schematic showing number of significant L-R pair interactions between olfactory sensory neurons and olfactory epithelium cells by STcomm analysis (top). The bottom panel showing the spatial mapping (color intensity) and colocalization of olfactory sensory neurons and olfactory epithelium cells according to deconvoluted weights in section 4 (S4). b. Dot plot showing significant L-R pairs between olfactory sensory neurons and olfactory epithelium cells via secreted signaling (left) and cell-cell contact (right) with p < 0.05. Color bar represents communication probability of L-Rs from TOME scRNA-seq data at E13.5. c. The spatial distribution of expression of L-R pairs of Wnt4 and Fzd3, and their co-expression in S4. The inlet shows the co-expression level of Wnt4 and Fzd3 in co-localized region of olfactory sensory neurons and olfactory epithelium. d. The spatial distribution of expression of L-R pairs of Efna5 and Epha4, and their co-expression in S4. e. Dot plot showing significant L-R pairs communication among spatial proximity brain related cell types calculated by STcomm. f. Schematic showing number of significant L-R interactions between neuron progenitor cells and inhibitory interneurons. g. Circos plots representing significant interaction of L-R pairs of Nrxn3 and Nlgn1 among spatial proximity brain related cell types. h. Spatial plots showing the spatial distribution (color intensity) and colocalization with spots of neuron progenitor cells and inhibitory interneurons according to deconvoluted weights in section 2 (S2). i. Spatial distribution of expression and co-expression of L-R *Nrxn3* and *Nlgn1* in S2. j. *Nrxn3* and *Nlgn1* spatial expression pattern examined by RNA-scope in brain tissue sections matched to S2 in (i). White dashed box showing the staining of *Nrxn3* and *Nlgn1* in spatial proximity cells. Blue, DAPI; green, Nrxn3; red, Nlgn1; yellow, co-location of Nrxn3 and Nlgn1.

Based on our spatial data, we obtained significantly co-expressed L-R pairs for spatially co-localized cell types by performing Fisher’s exact test ^51^ on binarized co-localized cell type pairs and co-expressed L-R pairs at spot level. We further calculated significant communication between L-R pairs in co-localized cell type pairs based on the reference single cell transcriptomic data and kept only spatially confident communications based on the above Fisher’s exact significance (Fig. 5c and Methods). Leveraging the spatial information, STcomm characterized confident cell-cell interaction within the tissue organization (Fig. 5c and Supplementary Table 8). According to the identified spatial co-occurrence cell types, we examined the LR interactions on olfactory epithelium and olfactory sensory neurons which showed high frequency interactions. We identified 53 L-R pairs from olfactory sensory neurons to olfactory epithelium and 69 L-R pairs from olfactory epithelium to olfactory sensory neurons (Fig. 6a). Notably, for secreted signalings, *Sema3* family, *Wnt* and *Bmp* sourced from olfactory epithelium showed a high communication probability to target olfactory sensory neurons (Fig. 6b), in line with their reported role in regulation of axon outgrowth and navigating of olfactory sensory neuron ^52–56^. Visualization of *Wnt4* and *Fzd3* expression on the tissues showed a high co-expression level in the co-occurrent spots of olfactory epithelium and olfactory sensory neurons (Fig. 6c). For cell-cell contact L-R between these cell-type pairs, *Efna-Epha* family and Notch signaling were identified with a strong communication probability, indicating their important role in regulating the development of olfactory epithelium cells ^57^ (Fig. 6d).

E13.5 embryogenesis is hallmarked by its rapid neurogenesis. We next focused on the specific interactions among neuronal cells that lay out the complex architecture composition (Extended Data Fig. 9a,b). We identified a set of L-R pairs with significant communication probability among these co-localized neuron related cells (Fig. 6e). Specifically, 23 L-R pairs source from neuron progenitors and 24 L-R pairs source from the inhibitory interneurons were identified between the neuron progenitors and the inhibitory interneurons (Fig. 6f). Among them, *Nrxn3*-*Nlgn1* showed prevailing communication probability between the neuron progenitors and the inhibitory interneurons (Fig. 6e,g and Extended Data Fig. 9c). Taken the brain-containing section 2 (S2) as an example, we found that the spatial co-occurrence of the neuron progenitor cells and inhibitory interneurons cell pairs overlapped well with the spatial co-expression of *Nrxn3*-*Nlgn1* LR pairs on tissues (Fig. 6h,i). Similar results were found in section 1 (Extended Data Fig. 9d). Furthermore, by examining the spatial expression of these L-R pairs using single-molecule fluorescent in situ hybridization ^58^, we confirmed the spatial proximity of *Nrxn3*-*Nlgn1* expression on brain tissue sections (Fig. 6j and Extended Data Fig. 9e), whereas the potential function of the *Nrxn3*-*Nlgn1* interaction in mediating neuron progenitor cells and inhibitory interneurons interaction awaits further exploration. Taken together, by integrating single cell transcriptome data, our spatial atlas efficiently decodes the spatial proximity cell-cell communications with STcomm.

## Discussion

Knowledge of organogenesis serves as the foundation for research in regenerative medicine, where generation of cells and tissues in vivo and in vitro commonly employ regulatory programs underlying organogenesis ^59^. Being anatomically organized and functionally distinct structure that is composed of various cell types, the spatial architecture of developing embryo is essential for functional display of normal development and homeostasis, as well as pathophysiology. Albeit our understanding of embryo organogenesis is rapidly accelerating, driven in particular by revolutionary single-cell transcriptomics, a key limitation of most single-cell molecular profiling methods is that they operate on disaggregated cells or nuclei so important spatial information is missing. With the advent of spatial transcriptomic technology, the natural state of cells in native tissues can be assayed to identify the location-defined cell types and understand how the cells are communicating with each other in setting up the body architecture.

While two-dimensional (2D) spatial transcriptome from a tissue slice has been informative to our understanding of the molecular organization of cell types in a static view, a stacked 3D spatially resolved transcriptomic analysis of aligned sections provides a dynamic angle to reveal the tissue architectures along the embryonic axes including A-P, D-V and L-R. In the current study, the systematic 3D spatial transcriptome from the head to the tail of mouse embryo organogenesis allows the construction of frameworks for organism-level molecular annotation based on 3D spatial coordinates. We further revealed the spatial gene expression profiles that determine the sophistical cellular structure of the E13.5 mouse embryo and we showed that the spatial heterogeneity of tissue organization is highly related to the organ compositions. We also uncovered critical and potential novel transcriptional regulators for tissue specific function development. Importantly, the 3D spatial profiling enables the identification of new genes of body axis along the anterior and posterior or dorsal and ventral, which greatly expands the insights for body patterning during embryogenesis. Leveraging the spatial information, we also delineated the confident L-R interactions with colocalized cell types across the embryo by STcomm.

Our study provided a high-quality resource for dissecting mouse organogenesis development and for developing new bioinformatics pipelines moving from 2D to 3D ^60^. In addition, the rich data of 3D spatial transcriptome of full embryos by transverse spatial data, provides new insights into the understanding of embryo development process and defining key genes for specific developing organ and early location. The spatially aware cell-cell interactions provide better views of cell location, communication and heterogeneity in a developing cell lineage than scRNA-seq per se. We also built an expandable web portal as spatial transcriptomic resource for further deciphering mouse organogenesis. We envision a compiling of more developmental stages on 3D spatial transcriptome dimension to generate a 4D atlas will greatly deepen our understanding of mammalian embryogenesis and cast insights for directed in vitro generation of various organs.

## Methods

### Embryo collection and Spatial transcriptome preparation

Mid-gestation organogenesis stage (embryonic day 13.5, E13.5) embryos were collected from C57BL/6 mice, and embryos images were taken for recording and confirmation of developmental staging. Animal procedures conducted in this study were approved by the Institutional Animal Care and Use Committee of Bioland Laboratory, Guangdong.

Collected embryos were embedded in Tissue freezing medium (OCT; Leica Microsystems, cat. no. 020108926) and stored at −80°C until sectioning. The whole embryo tissue was serially cryo-sectioned (Leica CM3050 S) along craniocaudal axis at 10 μm, and about 1000 sections were harvested. Considering the morphology and uniform sampling, the section 96, 189, 269, 361, 449, 541, 634, 732, 824 and 917 were selected for spatial transcriptomic analysis by modified 10x Genomics *Visium* platform. Briefly, the Visium Spatial Tissue Optimization Kit was used to optimize the permeabilization condition. The ideal embryo tissue permeabilization condition was set to 6 min. Selected sections were stained with 1% Cresyl violet solution and imaged using a Zeiss Axio Observer 7 microscope under a 10 lens magnification, then processed for spatial transcriptomics using Visium Spatial Gene Expression Kit (10x Genomics) according to the manufacturer’s instructions.

The resulting cDNA were synthesized, amplificated and then purified using AMpure beads. The cDNA library was assessed by Qubit 4.0 Fluorometer and Qsep100 Bio-Fragment Analyzer (Bioptic). The cDNA libraries were sequenced on Ilumina Novaseq 6000 system with paired-end 150 bp reads, aiming for 100 k raw reads per spots.

### In Situ Hybridization

Total RNA was prepared using GEO-seq extraction method ^61^ from the whole embryos of E13.5 and were further used as template for preparing probes. Whole embryo and spinal cord were successively fixed and dehydrated in accordance with Yoshihiro Komatsu et. al ^62^. Cryosections with 10 μm thickness were dried at room temperature for 10 minutes, then were fixed in 4 % paraformaldehyde (RNase free) and incubated for 15 minutes on ice. Slides were washed by PBS and incubated with 5 ug/ml proteinase K in PBS for 10 minutes at room temperature, and permeabilized in 2x SSC supplemented with 0.5% Triton for 30 minutes at room temperature. For pre-hybridization, slides were placed into the hybridization buffer (5xSSC pH4.5, 50% formamide, 0.1% Tween, 0.1% CHAPS,5 mM EDTA) at 65°C for 1 hour. After pre-hybridization, slides were hybridized with 500 ng/ml RNA probes in hybridization buffer (5xSSC pH4.5, 50% formamide, 1 mg/ml yeast RNA, 0.1 mg/ml heparin, 0.1% blocking reagent, 0.1% CHAPS and 5 mM EDTA) overnight at 65°C. Then, slides were successively washed by pre-hybridization buffer and 1:1 (pre-hybridization buffer and TNTE buffer(10mM Tris-HCl pH7.6,500mM NaCl,0.1% Tween 20, 1mM EDTA) for 30 minutes at 65°C,and washed again. After blocking, slides were incubated with Blocking buffer II containing 0.1% Anti-DIG-AP fab anti body (Roche, 11093274910) overnight at 4°C, and washed five times with buffer I for 15 minutes at room temperature, followed by washing twice with buffer III (100 mM Tris pH 9.5, 50 mM MgCl2, 100 mM NaCl) for 5 minutes. Finally, slides were incubated in BCIP/NBT Kit (Cwbio, CW0051S) for staining. The resultant stained slides were imaged with a CellCut laser microdissection system (CellCut System; MMI) microscope.

### RNAScope in situ hybridization

Fresh embryo was embedded in Tissue freezing medium(Leica) and stored at −80°C. For validation experiments, RNAscope Multiplex Fluorescent Reagent Kit v2 was used on fresh frozen embryo sections 10-μm thick from E13.5 C57BL mice in cryostat at −18°C (Leika CM3050 S), with Nlgn1 probes(Akoya biosciences 533511) and Opal 570 Reagent Pack (Akoya Biosciences ASOP570), Nrxn3 probes(Akoya biosciences 505431) and Opal520 Reagent Pack (Akoya biosciences ASOP520), Negative Control Probe (Akoya biosciences 320871), and Positive Control Probe (Akoya biosciences 320881) following the kit instructions. Images were acquired at 20× on an OLYMPUS VS200 microscope.

### Spatial RNA-seq data processing

#### Generation of spatial expression matrices

Quality control and adapter trimming on Raw reads were implemented with fastp-0.21.0 ^63^. Clean reads were mapped to mouse reference genome and gene annotations (mm10-3.0.0) using the Space Ranger v.1.0.0 (10x Genomics). To obtain only tissue-associated barcodes, spots were manually aligned to the tissue image with the Loupe Browser v.4.0.0 (10x Genomics). Count matrix were extracted by loading the output directory of Space Ranger into Seurat v.4.0.5 ^64,65^.

#### Data preprocessing

The expression datasets were filtered with cut-offs at a minimum of 1,000 detected genes and a maximum of 10% mitochondrial counts per spot. Because of the high quality of our spatial RNA-seq data, only less than 25 spots for each section were filtered.

All spot transcriptomes across 10 sections were merged together. The merged UMI counts was normalized by LogNormalize method with a default scale factor 10,000 and scaled by ScaleData function in Seurat with regressing out of sections and number of genes and counts per spot specified by the vars.to.regress argument.

#### Variable feature selection

For variable gene selection, we considered using both high variable genes (HVGs) and spatial variable genes (SVGs) by the following steps. First, we identified 2000 HVGs by vst method from FindVariableFeatures function in Seurat ^65^. Second, spatial variable genes were selected by two SVG identification methods. One is implemented through binSpect function in Giotto (R package, v1.0.3), a standard general-purpose toolbox for spatial transcriptomic data analysis, which includes a rich set of algorithms for characterizing tissue composition, spatial expression patterns, and cellular neighborhood and interactions^66^. The other is, implemented via SpatiallyVariableFeatures function with Trendseeck method in Seurat ^67^. Thus, Spatial variable genes were obtained by intersecting of genes identified by these two methods for each section. We then retrieved both HVGs and SVGs genes as variable gene for each section. At last, all variable genes in each section were combined together as the final variable gene list after filtering out mitochondrial and hemoglobin genes.

#### Dimension reduction, clustering and marker gene identification

Dimensionality reduction was performed with PCA and then top 50 PCs were used to create a shared NN graph and analyzed by Louvain clustering with a resolution of 1.2. UMAP project and visualization in 2D space were also applied on the above shared NN graph ^68,69^. Differential expressed marker genes for each cluster were identified by Wilcoxon rank sum test using FindMarkers and FindAllMarkers functions.

#### Gene Ontology enrichment and spatial domain annotation

We applied Metascape (http://metascape.org) ^70^, and clusterProfiler (R package, version 3.14.3) ^71^ to perform Gene Ontology (GO) enrichment analysis for each group of DEGs and regulon groups. In order to determine the identities of spatial domains, we combined unsupervised clustering of spatial expression profiles, spatial anatomy structure, enriched GO terms from differential expression genes and top marker genes with known literature curated cell type/region specific expression gene to annotate the major spatial domains.

### TF and Regulon analysis

To infer transcription factors and the gene regulatory network (regulon) in our spatial transcriptome data, we applied SCENIC (single-cell rEgulator Network Inference and Clustering) pipeline with pySCENIC (v0.10.3), a lightning-fast python implementation ^18^.

The procedure contains three main steps: 1) co-expression modules between TFs and the candidate target genes were first identified base on the correlation of normalized gene expression across all sample spots by GRNBoost2 with default parameter settings. Genes expressed in less than 10 spots were filtered. 2) co-expression modules were then further pruned by keeping only direct targets of TFs based on motif discovery by RcisTarget. Thus, modules composed of TF and TF direct target genes were defined as a regulon. 3) The Regulon Activity Score (RAS) for each spot was calculated through the area under the recovery curve by AUCell and the regulon activity for each spatial domain was computed as the mean activity of the corresponding spots.

### Calculation of regulon specific score (RSS) in spatial domains

We calculate the spatial domain specificity score of a regulon as describe in Suo et. al ^19^. Briefly, the RAS in all spots was normalized as a probability distribution and the indicator vector of a spot belongs to a specific spatial domain or not was also normalized as a probability distribution. Next the Jensen-Shannon Divergence (JSD) was computed between these two probability distributions. Finally, the RSS was calculated by converting the JSD to a similarity score. The selected regulons with top RSS for each spatial domain indicated their specificity and essentialness in the corresponding spatial domain.

### Regulon module analysis

To examine the co-regulation of TFs, we performed regulon modular analysis as described previously ^19^ which involves two steps. First, we calculate the Connection Specificity Index (CSI) for each pair of regulons following the instruction in ^29^, which is a context dependent metric used for identifying specific associating partners. Next, regulon modules were identified by clustering the regulons with hierarchical clustering based on Euclidean distance of the CSI matrix.

To study the relationship among different regulons, we also build the regulon co-activation network based on CSI matrix with a threshold of 0.85 to filter weak connected regulons and visualized the network by cytoscape (v3.9.1) ^72^. The regulon module activity score of each spatial domain was calculated by averaging the activity scores of all regulon members in spots which belongs to the corresponding spatial domain. For each module, the top correlated spatial domain was identified according to the regulon module activity score.

### Spatial pattern of spinal cord analysis

To systematically exploring the spatial patterning from anterior-posterior (A-P) axis, we conducted the pseudo-space analysis in the spots from spatial domain of hindbrain and spinal cord. All Hox genes expressed in the dataset were used for reconstructing the pseudo-axis by Monocle 2^73^. The expression of typical A-P patterned Hox genes were smoothed in extracted spots which were first ordered along the sections from anterior to posterior and then ordered by predicted pseudo-space within each section.

To identify A-P related genes beside Hox genes, we applied the following steps: 1) identified significant differential genes among the A-P slices by differentialGeneTest function in Monocle 2 with the pseudo-time formula. 2) Pearson correlation between differential expressed genes and combinatorial patterns of A-P slices were calculated. The union of genes with top 20 Pearson Correlation Coefficient for each combinatorial pattern were selected as strong AP-related genes.

To examine the spatial patterning along dorsal-ventral axis, we retrieved a list of marker genes which were used to identify D-V domains of neuronal progenitors and neuron clusters from Delile et al ^41^. As our 10x Visium spatial transcriptomics are not in single cell resolution, we separated the predefined dorsal ventral marker genes into Dorsal (D), media (M) and ventral (V) domain related gene sets for both the inner progenitors and the outer neuron regions. In brief, for neuronal regions, D marker gene sets were defined by combinatorial markers of dl1-dl6, M marked by V0-V2b and V marked by Mn and V3. Similarly, for neuron progenitor regions, D marker gene sets include combinatorial markers of RP and dp1-dp6, M includes markers of p0-p2, V includes pMN, p3 and FP maker genes. Then, the activity score of each region related gene set was calculated by AUCell (v1.8.0) ^18^. For visualization, the activity scores were scaled and scores lower than the binary assignment score were set to 0 in outer neuron region. In progenitor region, the z-score lower than 2.5 were set to 0.

To identify novel D-V related genes, we divided the spinal cord region into dorsal, middle and ventral domains according to the D, M, V region related activity score. Differential expression analysis was conducted among these three regions by FindallMarkers Function in Seurat. Significant DEGs with adjust p-value less than 0.05 were considered as D-V related genes. For neuronal progenitor genes identification, we calculated differential expressed genes by comparing D17-ependyma and D16- medulla oblongata and spinal cord on sections from S4 to S10 based on the union of the marker genes of D16 and D17.

### Signaling pathway activity analysis in spatial transcriptomic data

Signaling activity scores were computed using the *AddModuleScore* function from Seurat based on the literature curated critical development related signaling signatures, including BMP, Wnt, Nodal, Fgf, Hedgehog, Hippo and Notch signaling pathway genes. Briefly, the score of each spot was computed as the average expression of each signaling signature subtracted by the aggregated expression of random control gene sets ^74^.

Cell cycle phase activity scores were calculated for all spots by CellCycleScoring function from Seurat function with mouse cell cycle-related genes retrieved from Giladi et al. ^75^.

### Spatial mapping of single cell types of mouse organogenesis atlas

To spatially mapping cell types of mouse organogenesis, we used the single-cell dataset of Trajectories Of Mammalian Embryogenesis (TOME, downloaded from http://tome.gs.washington.edu/) at the E13.5 stage as reference ^1^, and performed deconvolution for each section by the robust cell-type decomposition (RCTD, v1.0.4), which is a supervised learning method to accurately decompose the spatial transcriptomic mixtures for each pixels by using a scRNA-seq reference containing cell-type classifications (Cable et al., 2022). Before running the RCTD, we excluded Hemoglobin and Mitochondrial genes from both single-cell and our spatial transcriptome datasets. The method of multi-mode was selected to perform deconvolution analysis on our spatial data with default parameters, except for CELL_MIN_INSTANCE = 25, UMI_max = 2e+08. We extracted the cell type deconvolution weights of each spot for downstream analysis and visualization.

### Cell type co-localization within spots and cell-cell communication analysis

Spot-wise Pearson Correlation Coefficient (PCC) of weights generated from RCTD deconvolution results were calculated. Cell type colocalization network was created base on the PCC filtered with a threshold of 0.06 and adjust p-value less than 0.05 ^77^. The remaining cell-type connections were visualized by Cytoscape ^72^.

To study cell type interactions in spatial microenvironment, we develop an analysis workflow named STcomm (Fig. 5c). It combined the spatial cellular colocalization of single cells and ligand-receptor co-expression from spatial transcriptomic data and cell-cell communication based on the expression of ligand receptor pairs in single cell transcriptome. Firstly, we calculated co-localized cell types pairs and performed binarization according to the decomposed cell-type weights. Secondly, we extracted the expression of each pair of LRs in our spatial atlas, and obtained the binarized matrix of the spatial co-expressed LR pairs according to whether L(exp)*R(exp)>0. The ligand-receptor information was extracted from the CellChatDB.mouse database ^78^. Thirdly, Fisher’s exact test was performed on the cell types co-localization matrix and LRs co-expression matrix obtained above, and spatially co-expressed LR pairs in the co-localized cell type pairs were identified according to adjusted p-value < 0.05. At last, we ran cell-cell communication by CellChat (v1.4.0) with the default parameters on the dataset of E13.5 TOME and kept the communication results with the above identified significant spatially co-expressed LR pairs within co-localized cell type pairs .

### Web service

The Mouse Organogenesis Spatial Transcriptomic dataset can be interactively explored at our website, which was constructed to navigate the spatial atlas of all 10 sections from E13.5 embryo using Dash in python. This web service provided five parts for data exploring, to search for the spatial resolved gene expression by the ‘Spatial Transcriptomics Explorer’, to identify the spatial regulon modules on each section by the ‘Regulation’, to explore the clustering and annotation of spatial domain based on molecular signatures by the ‘Spatial Domain Explorer’, to retrieve the spatial pattern of co-expressed genes by the ‘Gene Pattern Search’, and to display the 3D reconstruction of spatial domain of 10 sections from E13.5 embryo either illustrated by package plotly in Python or ImageJ. To reconstruct the 3D embryo model, the images of TS22 corresponding to stage of E13.5 were retrieved from eMouse atlas (www.emouseatlas.org), we then registered our experimental images manually with the matched images in TS22, and stacked all these images together by ImageJ.

## Acknowledgments

This work was supported in part by National Key R&D Program of China (2018YFA0801402), the “Strategic Priority Research Program” of the Chinese Academy of Sciences (XDA16020404), National Natural Science Foundation of China (32270854, 32161160322, 31871456 and 32100483), Guangdong Basic and Applied Basic Research Foundation (2019B151502054, 2019A1515110985 and 2020A1515110517), Science and Technology Program of Guangzhou (202102080293), Frontier Research Program of Bioland Laboratory (Guangzhou Regenerative Medicine and Health Guangdong Laboratory, 2018GZR110105013), Jiazi Research Innovative Project of Bioland Laboratory (2019GZR110108001), Science and Technology Planning Project of Guangdong Province (2020B1212060052). We thank XL. Peng, XZ. Zhai and LF. Liu for experimental support. We thank P. Tam, N.Sheng, G.Bai, N. Jing for discussion and critical reading of this study.

## Author Contributions

G.Peng, and F.Qu designed the study. G.Peng, S.Suo, G.Cui and F.Qu supervised the project. F.Qu, W.Li, X.Ren, F.Lu, and R.Zhang. analyzed the data with contributions from J.Ke and Z.Zhang. J.Xu, G.Cui. and X.Meng performed the experiments with contributions from L.Qin, J.Zhang, X.Luo X. Zhou and M. Wang. G.Wu, J.Chen and D.Pei provide important reagents and suggestions. G.P., and F.Q. wrote the paper with the help of W.Li, G.Cui, S.Suo and J.Xu.

## Data availability

The mouse organogenesis spatial transcriptomic data generated in this study are available under NGDC and can be explored at the web portal. All other data are available from the corresponding authors upon request.

## Code availability

Source code for STcomm is available at https://github.com/gpenglab/STcomm.

