## Supplemental information for "Three-dimensional molecular architecture of mouse organogenesis"

### Supplementary Inventory

1.Extended Data Fig. 1 to 9

2.Supplmentary Tables 1 to 8

### Supplementary Figure Legends

**Extended Data Fig.1| Data Quality and replicates of spatial transcriptional atlas for mouse embryo development in organogenesis at E13.5.**

(a) Distribution of UMI count (upper), genes (middle) and mitochondrial percentage (bottom) for the collected 10 sections.

(b) The total number of spots in each tissue section included for downstream analyses.

(c) Pearson correlation coefficient (R=0.986) of pseudo-bulk profiles for section 1 (S1) from mouse embryo 1 (E1) and similar section of embryo 2 (E2).

(d) Pearson correlation coefficient (R= 0.991) of pseudo-bulk profiles for section 2 (S2) from E1 and similar section from E2.

(e,f) UMAP embedding of spots from S1 of E1 and E2, colored by sample identities (e) and clustering of spatial regions (f). Colors represent cluster assignment based on Louvain clustering of all spots from the two sections.

(g,h) 5 clustered spatial regions showed on tissue section1 of E1 (g) and E2 (h). The same color scheme as (f) for spot colors.

(i)Heatmap showing pairwise spatial cluster correlations of section1 of E1 and E2.

(j,k) UMAP embedding of spots from S2 of E1 and E2, colored by sample identities (j) and clustering of spatial regions (k).

(l,m) 6 clustered spatial regions showed on tissue section2 of E1 (l) and E2 (m).

(n)Heatmap showing pairwise spatial cluster correlations of section2 of E1 and E2.

(O) Spatial visualization of expression for selected marker genes (*Ina*, *Col1a1*, *Foxg1*, *Lhx2* and *Dbx1*) of the S1 and S2 from two embryos (E1 and E2) and ISH images of the related genes from MGI database and Allen Brain Atlas.

**Extended Data Fig.2| Spatial domains and the signature genes at E13.5.**

(a) Stacked violin plots showing the expression levels of the specific markers in each of the spatial domain.

(b) Highlighted spatial domains mapping across all the ten sections (left) and the spatial expression of specific marker genes for each spatial domain across all the ten sections (right).

(c) Percentage of spatial domains assigned in each section.

**Extended Data Fig.3| Gene regulation network in the spatial domains.**

(a,b) UMAP embedding of all spots based on regulon activity scores (RAS). Colors represent spatial domain assignment based on gene expressions (a) or RAS (b).

(c) Concordance between spatial domains based on gene expression and RAS. The color bar of heatmap represents the percentage of spots in spatial domains (rows) that was labeled as the indicated regulon clusters (each row of proportions was summed to 1).

(d) Spatial distribution of RAS on tissues for selected specific regulons and related TFs in the spatial domains.

**Extended Data Fig.4| Spatial transcriptomic annotation of visceral organs at E13.5.**

(a) Scatter plots showing the statistical summary of differential expressed genes in each subclusters. X axis indicates the log2 foldchange of average expression between given spatial subcluster and all others in D5 domain, Y axis and the size of dots indicates the difference of percentage of gene expression detected between given subclusters and all others, and colored by -log10(p-adjust). Top different genes were indicated.

(b) Spatial distribution of visceral organ subclusters and corresponding spatial expression of selected marker genes on different tissues.

(c) Enriched GO terms of differential marker gene in each subcluster. Color bar indicated the adjusted p-value and dots are scaled by the gene ratio.

(d) Rank of regulons for lung based on regulon specificity score (RSS) (left) and spatial visualization of the activity score of top regulons Nkx2-1and Tbx4 (right).

(e) Rank of regulons for gut based on RSS (left) and spatial visualization of the Activity Score of top regulons Cdx2 and Isx (right).

**Extended Data Fig.5| Spatial patterning for spinal cord in mouse embryo organogenesis at E13.5.**

(a) Heatmap showing the expression pattern of the identified A-P axis related genes in hindbrain and spinal cord from section 2 to section 10 along anterior to posterior.

(b) Spatial expression of *Pou4f1* in the hindbrain and spinal cord tissue spots across section 2 to 10 (left) and ISH validation of *Pou4f1* expression in the spinal cord (middle) and whole embryo (right). Arrows denote the D-V or A-P localization.

(c) Heatmap of differentially expressed genes in Dorsal (D), Media (M), Ventral (V) regions of spinal cord along D-V axis.

(d) Spatial expression of *Ttyh1* in the hindbrain and spinal cord tissue spots across section 2 to 10 (left), and ISH validation of *Ttyh1* expression in medial region of spinal cord along radial axis.

**Extended Data Fig.6| Spatial mapping of cell types from mouse organogenesis at E13.5.**

(a) Bar plot showing the proportion of cell types occupied in spots across all embryo tissue sections after mapping back to the spatial regions.

(b) Spatial visualization of deconvoluted weights of 16 cell types.

**Extended Data Fig.7| Cell type compositions in spatial domains across the embryo.**

(a) Spatial visualization of deconvoluted weights of neuron related cell types after spatial mapping.

(c) Spatial visualization of deconvoluted weights of posterior floor plate cells and the expression of marker genes (*Slit1*, *Ntn1* and *Shh*) of posterior floor plate cells from TOME dataset (Qiu et al., 2022) across all embryo tissue sections.

**Extended Data Fig.8| Cell type compositions in spatial domains across the embryo.**

(a) Spatial visualization of deconvoluted weights of definitive erythroid cells, liver endothelium, megakaryocytes, amniochorionic mesoderm A and brain related cell type after spatial mapping.

(b) The top five abundant cell types that localized in each spatial domain. The cell types were selected by the deconvoluted weights greater than 0.05.

**Extended Data Fig.9| Cell-cell communication based on spatial and TOME single cell RNA-seq data.**

(a) Network plot showing the colocalized cell type pairs. The edge width is proportional to the indicated PCC of the connected cell type pairs. The orange color indicated the brain related cell types, and the green colors indicated all other cell types.

(b) The interaction network of significant L-Rs between pair of two cell populations from spots co-occurred brain related cell types by STcomm. The edge width is proportional to the indicated number of L-Rs.

(c) Dot plots showing the expression distribution of *Nrxn3* and *Nlgn1* at different spatial domains in our ST data (left) and at different cell populations in TOME single-cell data at E13.5 (right). The dot color and size represent the average expression and percentage of spots in each group.

(d) Spatial plots showing the spatial distribution (color intensity) and colocalization of neuron progenitor cells and inhibitory interneurons according to predicted weights by deconvolution in S1 (Top panel). The bottom plots showing the spatial distribution of expression and co-expression of LRs Nrxn3 and Nlgn1 in S1, which is similar to Fig.6i. but illustrated on the embryo tissue of section 1.

(e) *Nrxn3* and *Nlgn1* spatial expression pattern detected by RNA-scope in brain tissue section matched to section 1 in (d). White dashed box showing the staining of *Nrxn3* and *Nlgn1* in spatial proximity cells.

**Supplementary Tables**

**Supplementary Table 1**: The differentially expressed genes identified in all major spatial domains.

**Supplementary Table 2:** The ranked regulon specific score (RSS) in all major spatial domains.

**Supplementary Table 3:** The differentially expressed genes identified in 10 subclusters of spatial domain 5-visceral organ with smooth muscle.

**Supplementary Table 4:** The regulon specific score (RSS) of top 5 regulons in 10 subclusters of spatial domain 5-visceral organ with smooth muscle.

**Supplementary Table 5:** The top Anterior-Posterior axis related genes identified in spinal cord region.

**Supplementary Table 6:** The differentially expressed genes identified in Dorsal, Medial and Ventral region of spinal cord.

**Supplementary Table 7:**The predicted deconvolution weight (proportion) of confident cell types for each spot pixel by RCTD on multi-mode.

**Supplementary Table 8:** The spatial resolved cell-cell communication by integrating the spatial atlas and single-cell data with STcomm.

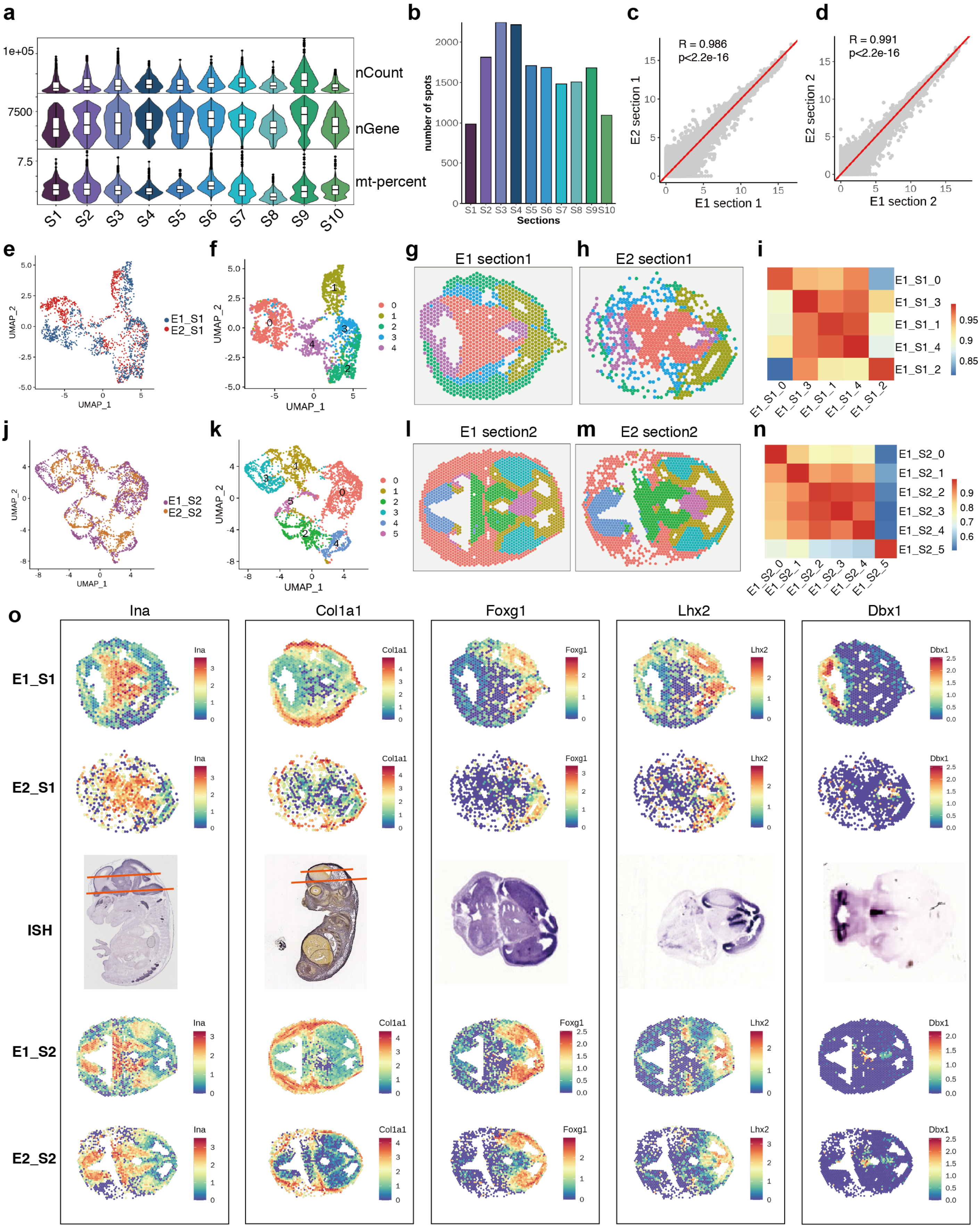


Extended Data Fig. 1

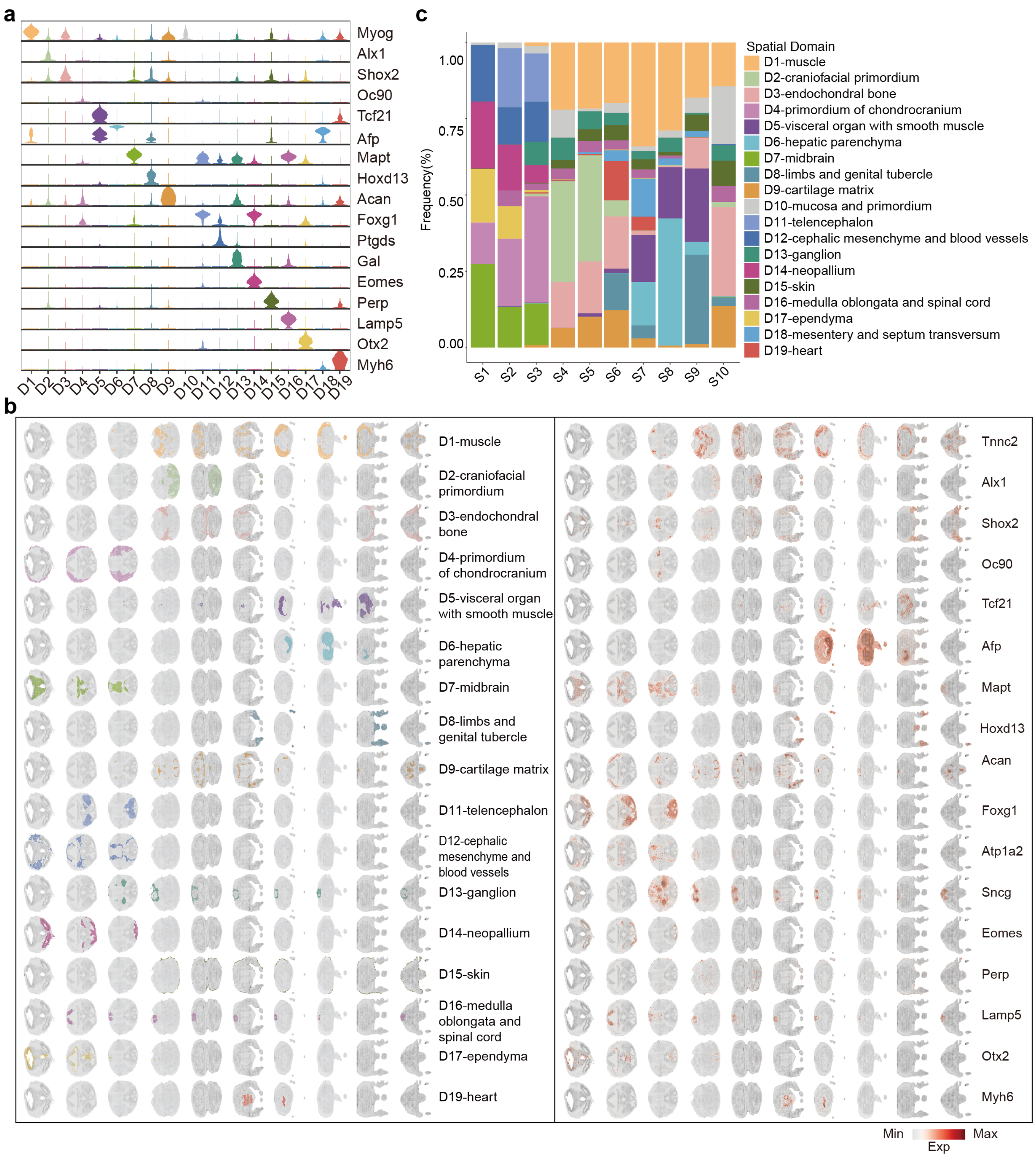


Extended Data Fig. 2

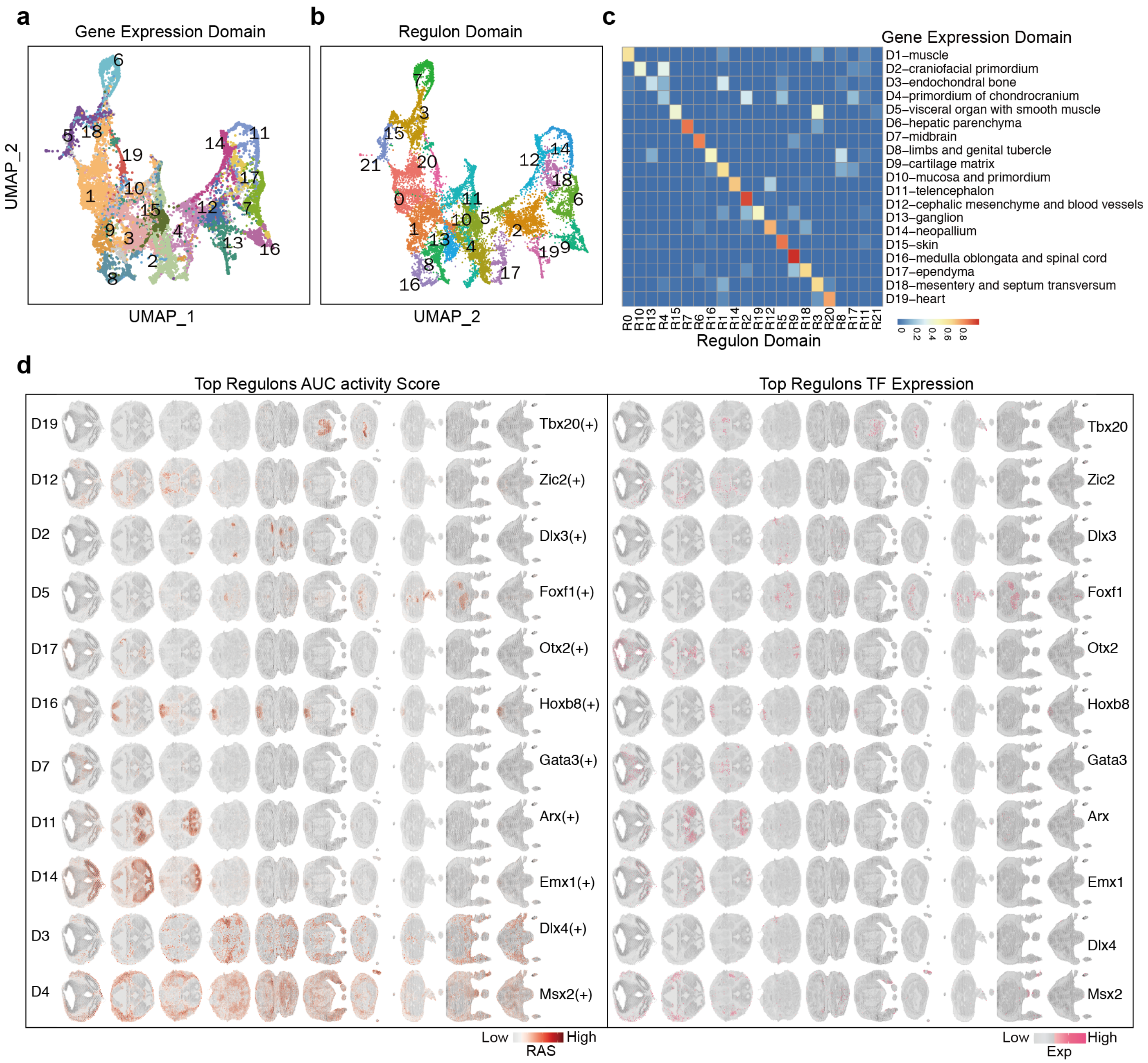


Extended Data Fig. 3

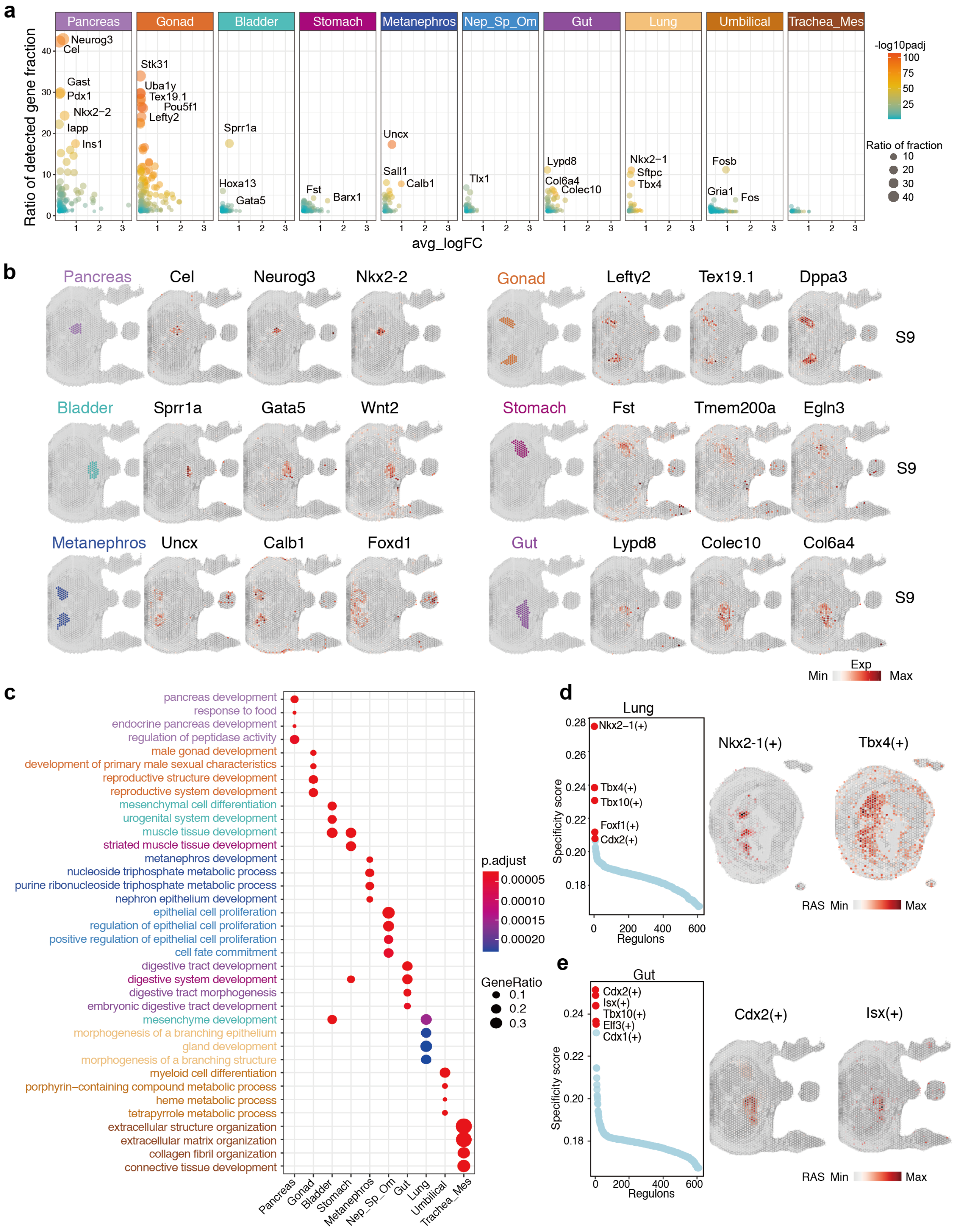


Extended Data Fig. 4

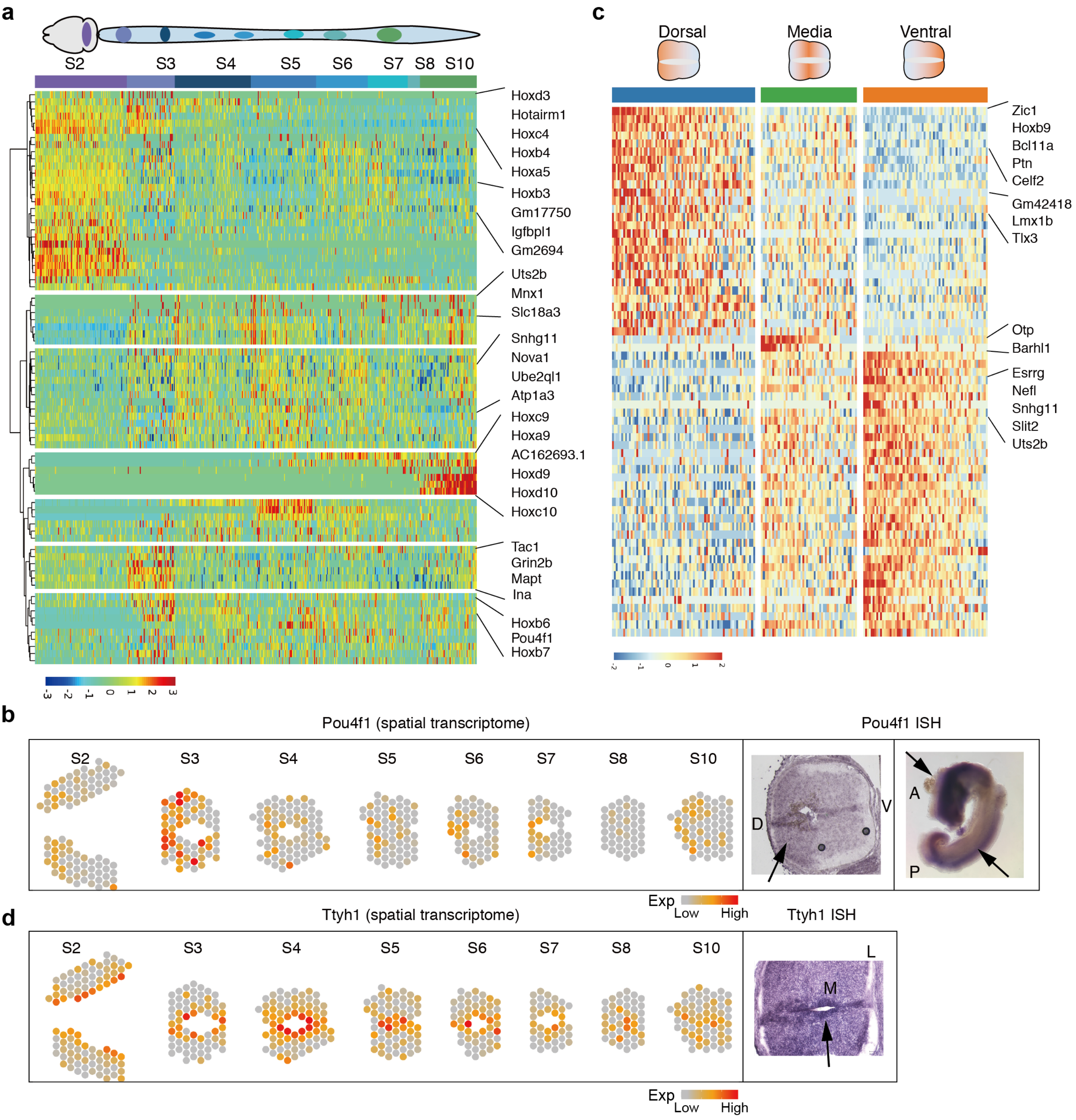


Extended Data Fig. 5

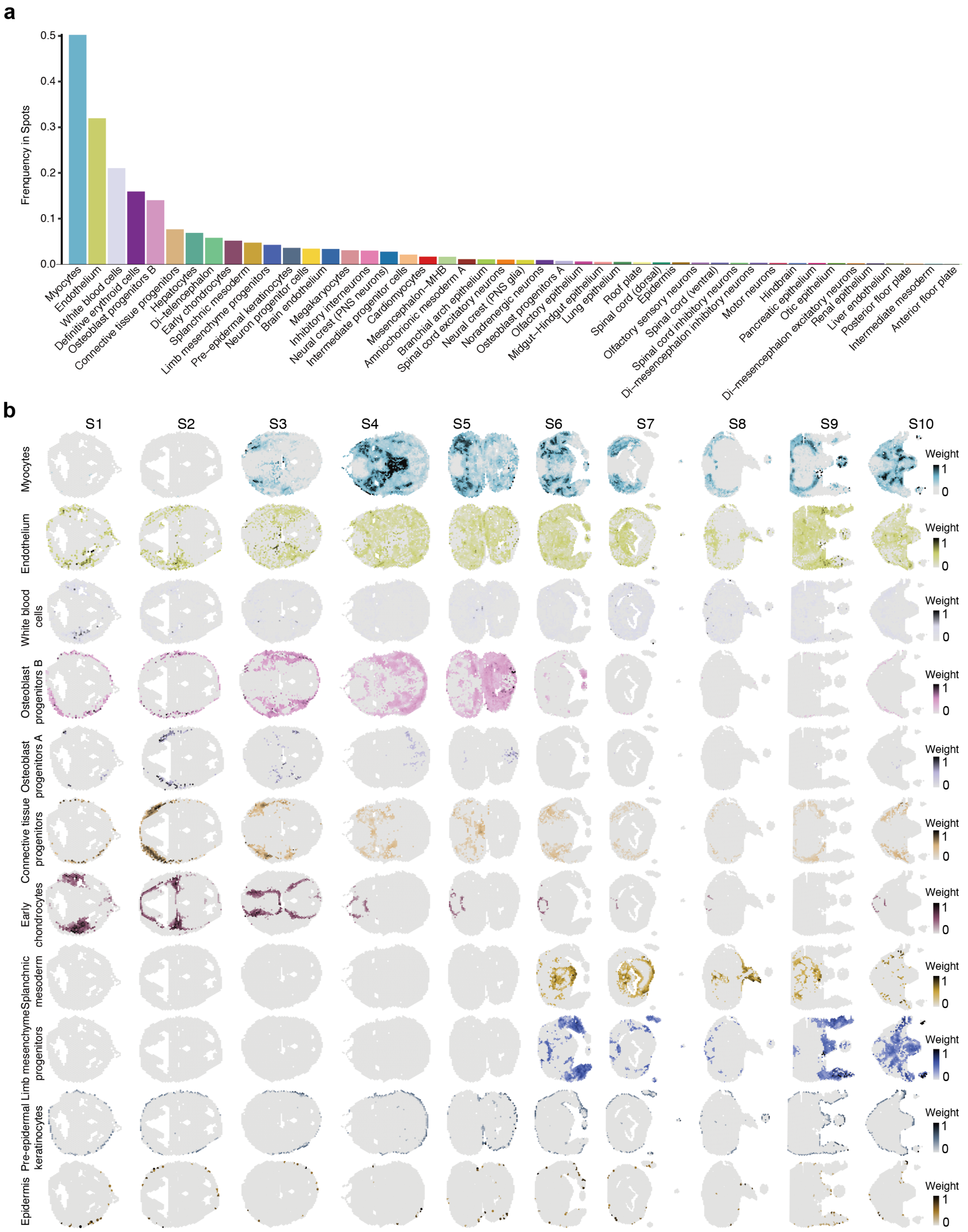


Extended Data Fig. 6

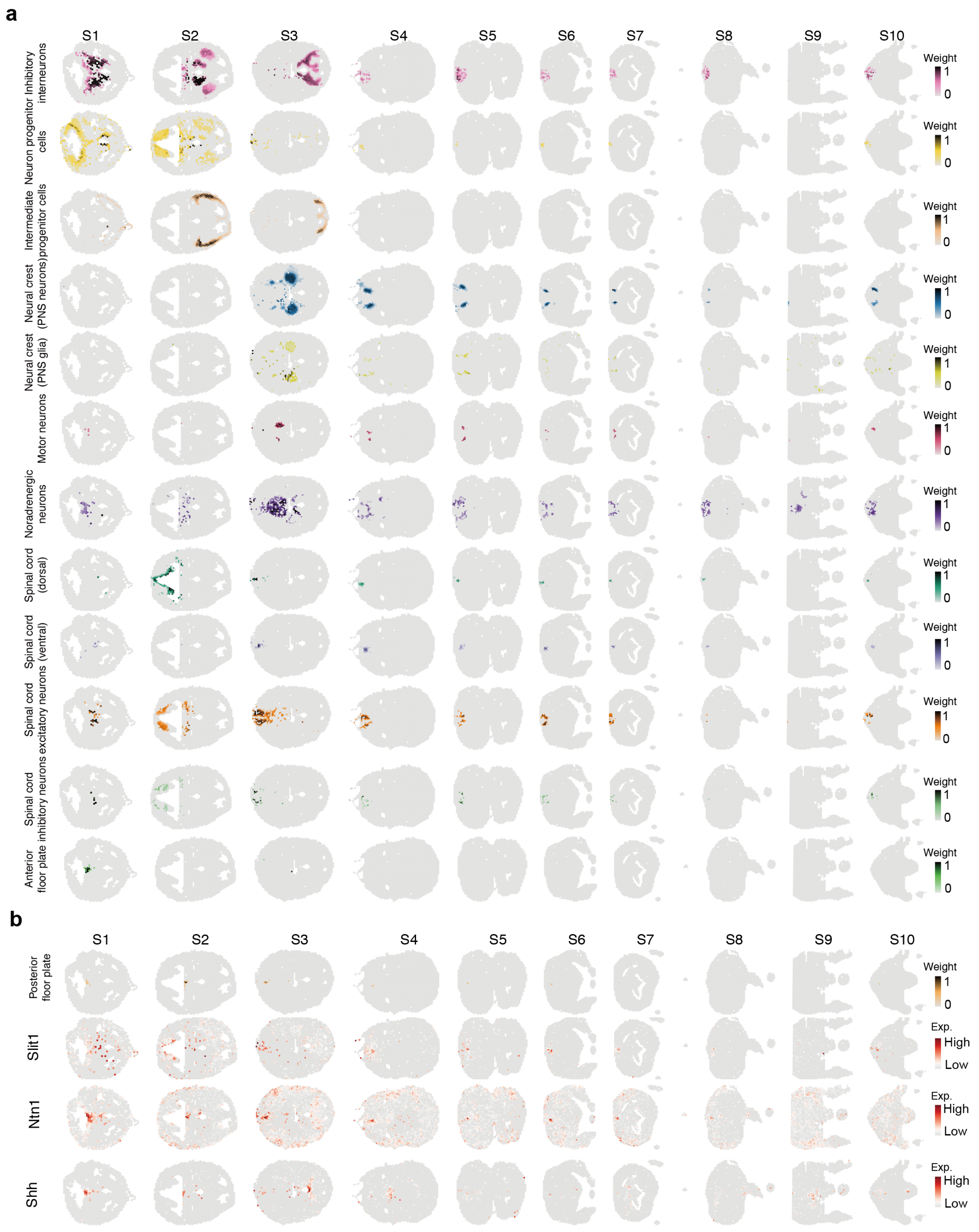


Extended Data Fig. 7

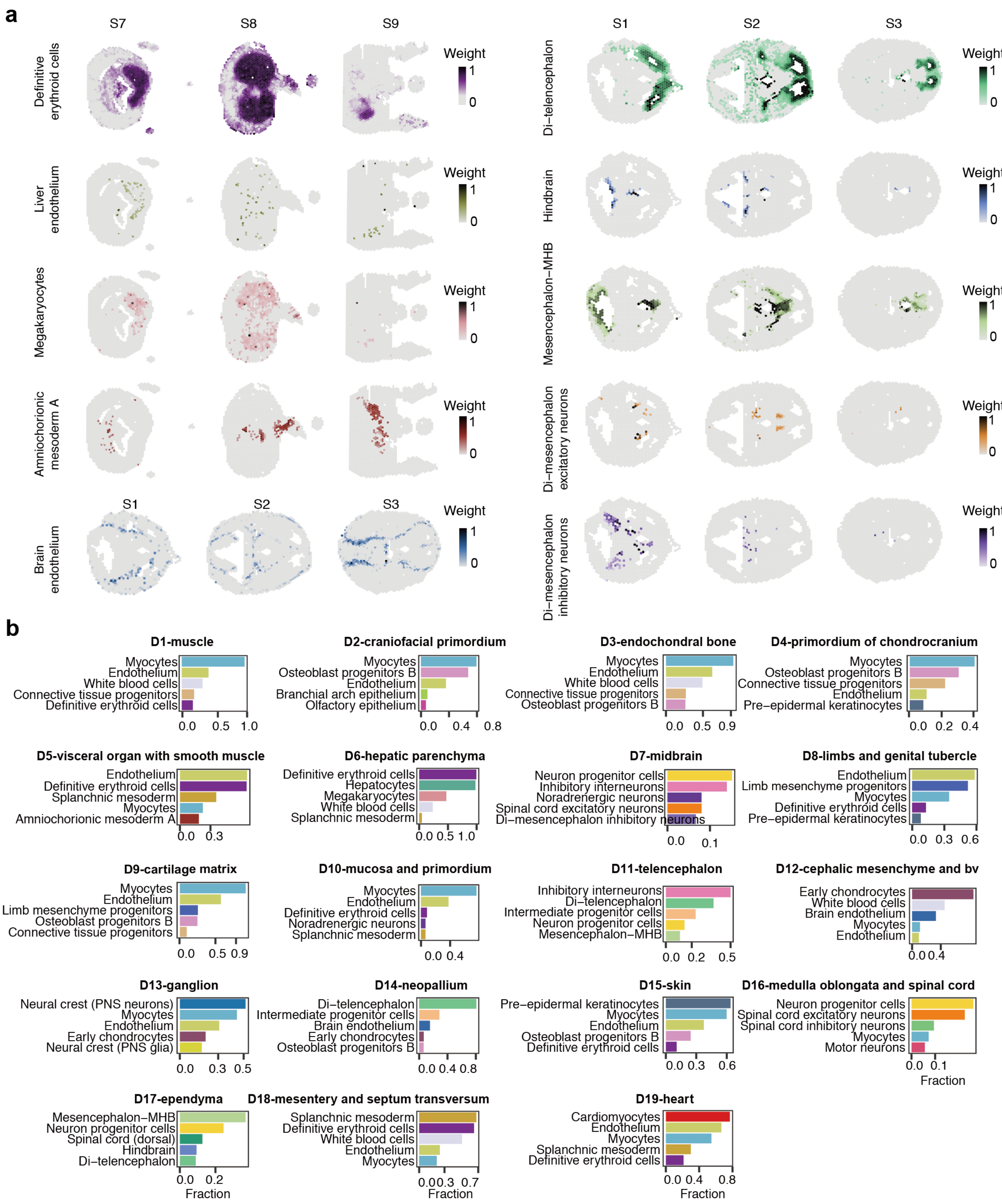


Extended Data Fig. 8

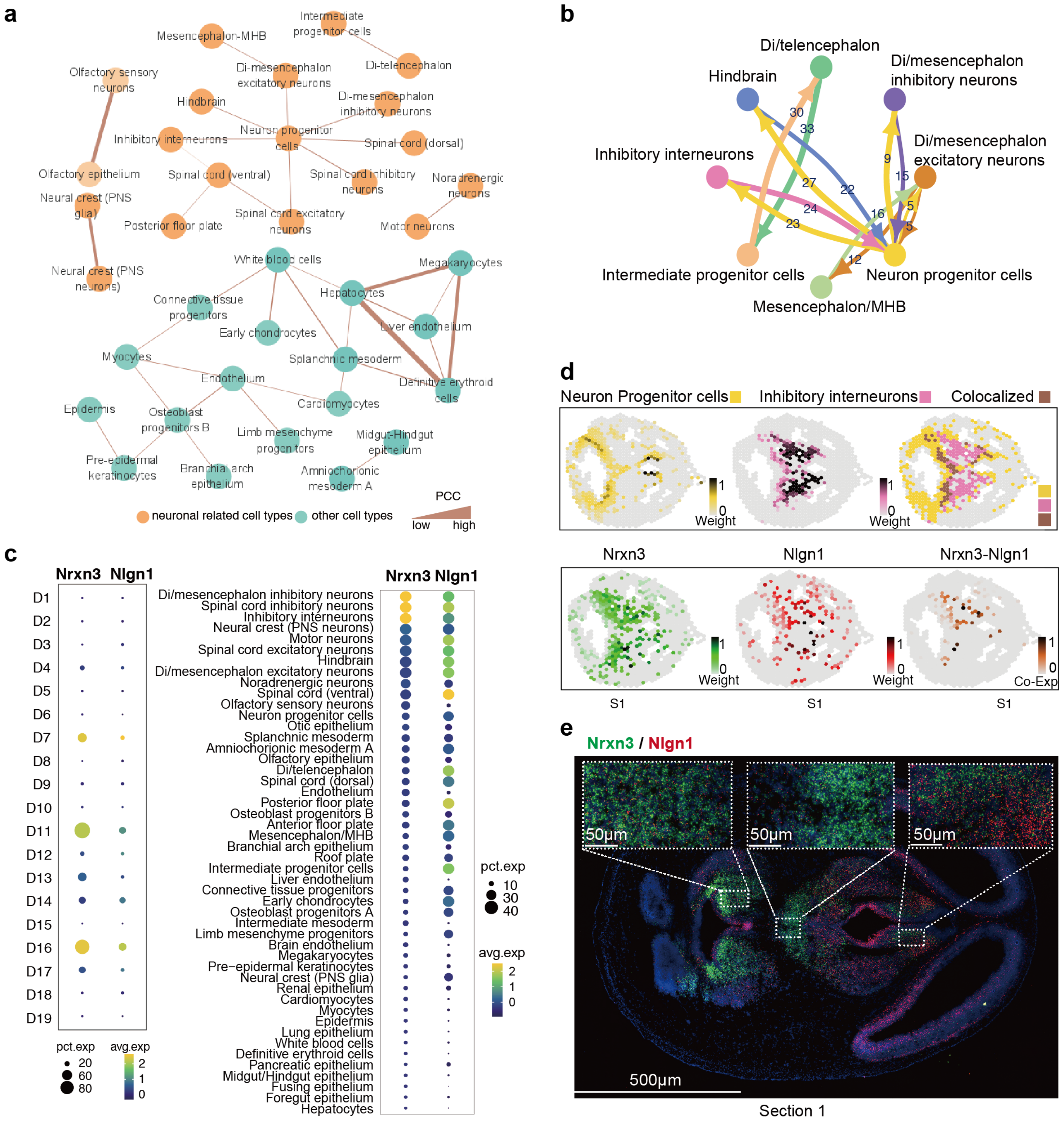


Extended Data Fig. 9
